# Intraplacental injection of human iPSC-derived PDX1+ pancreatic progenitors prolongs Pdx1-deficient mice survival

**DOI:** 10.1101/2024.05.09.593461

**Authors:** Arata Wakimoto, Zeynab Javanfekr Shahri, Hyojung Jeon, Takuto Hayashi, Ching-Wei Liao, Natalia Gogoleva, Fabian Suchy, Atsushi Noda, Yuri An, Hiromitsu Nakauchi, Yohei Hayashi, Michito Hamada, Satoru Takahashi

**Author notes:** These authors contributed equally to this work and may list their name first in their curriculum vitae (Arata Wakimoto, Zeynab Javanfekr Shahri). Correspondence (Y.H.), (M.H.), (S.T.).

## Abstract

Interspecies chimeras comprising human tissues have potential for use in disease modeling and regenerative medicine. Here, we successfully transplanted human induced pluripotent stem cell (iPSC)-derived PDX1+ pancreatic progenitor cells into *Pdx1*-deficient mouse embryos via intraplacental injection. The engrafted human cells predominantly localized to the duodenum, produced insulin, and extended the lifespan of *Pdx1*^-/-^ mice by up to 10 days after birth. Transcriptomic analyses confirmed human pancreatic gene expression in human cells engrafted into the mouse duodenum. Our findings demonstrated the feasibility of generating interspecies chimeras with functional human pancreatic cells through *in utero* transplantation of lineage-committed progenitors. This approach circumvents developmental barriers while minimizing ethical concerns associated with PSCs. However, the incomplete rescue of the *Pdx1*^-/-^ phenotype highlights the need for further research to enhance human cell engraftment and tissue integration. Overall, this study provides a foundation for developing human-animal chimera models for studying human development and regenerative therapies.

## Introduction

Interspecies chimeras with human tissues represent a promising approach for developing human disease models and generating organs for transplantation. Successful intraspecies (mouse-mouse) (Hamanaka et al., 2018) and interspecies (mouse-rat) (Goto et al., 2019; Kobayashi et al., 2021, 2010; Wu et al., 2017) chimeras have been established through blastocyst complementation. However, integrating functional human tissues into animal models has proven challenging (Hu et al., 2020; Maeng et al., 2021; Masaki et al., 2015; Wang et al., 2024).

An alternative approach for generating chimeric animals is to inject tissue stem/progenitor cells into a developing embryo using *in utero* injections (Cohen et al., 2020; Fleischman and Mintz, 1979; Fujiki et al., 2003; Jeon et al., 2021). Previous studies using neural crest cells successfully engrafted human cells into mouse embryos (Cohen et al., 2020, 2016). Utilizing lineage-committed cells minimizes the likelihood of donor cell differentiation for unintended purposes such as germline transmission and carcinogenesis (Crane et al., 2019; Wu et al., 2016).

Human pancreatic xenograft models are generated using newborn and adult NSG mice (Augsornworawat et al., 2020; Ma et al., 2018). These models have provided valuable insights into the functionality of human induced pluripotent stem cell (iPSC)-derived pancreatic cells *in vivo*. However, successful transplantation of human pancreatic progenitors into mouse embryos has not yet been accomplished because of the low survival rate of *in utero* injection systems.

Previously, we developed an intraplacental injection method to deliver cells (Jeon et al., 2021) and virus vectors (Gogoleva et al., 2023) into mouse post-implantation embryos through the placenta. This technique offers higher embryonic survival rates (>80%) compared to direct *in utero* injection methods because it avoids direct embryonic penetration.

Along with the placental connection established at E9.5, the initial indication of pancreas formation occurs as the dorsal endodermal epithelium thickens just below the stomach (Puri and Hebrok, 2007). All known pancreatic cell lineages originate from a small cluster of precursor cells within the foregut endoderm as defined by the expression of pancreatic and duodenal homeobox 1 (*Pdx1*) (Byrnes et al., 2018). *Pdx1* is recognized as a master regulator of pancreatic development, and its deficiency results in a lack of the whole pancreas and postnatal lethality within a day (Offield et al., 1996).

Various methods exist for differentiating iPSCs into pancreatic cell types, especially pancreatic β cells (Liu et al., 2014; Loh et al., 2014; Ma et al., 2018; Pagliuca et al., 2014; Rezania et al., 2014; Shahjalal et al., 2014). Most differentiation protocols follow a stepwise approach involving the progression of PSCs to definitive endoderm, pancreatic progenitors, and finally, functional pancreatic β cells.

Therefore, we hypothesized that introducing human iPSC-derived PDX1+ pancreatic cells into *Pdx1*-deficient mouse embryos would enable the derivation of human pancreatic tissue by occupying this vacant developmental niche. In this study, we investigated a method to generate a mouse model with human pancreatic tissue by transplanting iPSC-derived human PDX1-positive cells into *Pdx1*^-/-^ mice lacking a pancreas. The resulting human-mouse chimera exhibited prolonged survival compared with that of uninjected *Pdx1*^-/-^ mice. The engrafted donor-derived cells were capable of secreting human insulin. Notably, the injected human cells were primarily engrafted ectopically, predominantly in the duodenum. Our findings suggest that the *in utero* transplantation of lineage-committed human cells can give rise to interspecific chimeras with functional human-derived cells.

## Results

### Establishment and characterization of engineered human iPSC lines

To conduct the intraplacental injection of human PDX1-expressing cells into *Pdx1*^-/-^ mouse embryos, we first established two novel human iPSC lines with different tracking capabilities using CRISPR-mediated genome editing. The PDX1-P2A-tdTomato reporter line was generated by inserting a P2A-tdTomato cassette at the endogenous *PDX1* locus, enabling direct visualization of PDX1 expression during differentiation. The EEF1A1-iRFP670 line was created by integrating near-infrared fluorescent protein iRFP670 under the constitutive EEF1A1 promoter for robust cell tracking after injection. Both lines underwent characterization to confirm their pluripotent properties and differentiation potential.

Immunostaining revealed robust expression of core pluripotency markers OCT3/4 and NANOG in undifferentiated colonies of both PDX1-tdTomato and EEF1A1-iRFP670 lines (Figure S1A, C). To assess differentiation potential, we performed embryoid body formation assays followed by immunostaining for lineage-specific markers. Both lines successfully differentiated into all three germ layers, as evidenced by expression of ectoderm markers (PAX6, TUJ1), mesoderm markers (HAND1, SMA), and endoderm markers (SOX17, AFP) (Figure S1B, D). These results confirmed that our newly established lines retained the fundamental pluripotent characteristics for subsequent pancreatic differentiation studies.

### Pancreatic progenitor differentiation protocol optimization and validation

We adapted established protocols (Pagliuca et al., 2014; Rezania et al., 2014) for stepwise pancreatic differentiation, involving initial definitive endoderm induction with Activin A and CHIR99021, followed by pancreatic specification using retinoic acid, cyclopamine-KAAD, SB431542, and DMH-1 (Figure 1A). Using the PDX1-tdTomato reporter line, we confirmed 66.3 ± 7.4% colocalization between endogenous PDX1 protein and tdTomato expression at day 13 (Figure 1B). Flow cytometric analysis revealed that approximately 90% of viable cells expressed PDX1-tdTomato at day 13 (Figure 1C). We monitored the expression of key developmental markers by reverse transcription-quantitative PCR (RT-qPCR). From days 3 to 5, the definitive endoderm (DE) marker *SOX17* was upregulated (log2 fold change = 13). Subsequently, the pancreatic progenitor (PP) marker *PDX1* expression increased significantly (log2 fold change = 7), followed by expression of *NKX6-1* (log2 fold change = 4) by day 13 (Figure 1D). We validated our protocol using additional genetically independent human iPSC lines, WTC11 (GM25256, Coriell Institute), and NB1RGB (HPS5067, RIKEN Cell Bank). RT-qPCR analysis revealed consistent temporal dynamics of *PDX1*, *SOX17*, *HNF6*, *NANOG* across all three lines throughout the differentiation (Figure S2A-B). PDX1 immunostaining at day 13 showed PDX1-positive cells for the three lines (Figure S2C). These results established robust pancreatic progenitor differentiation across genetically distinct backgrounds. Furthermore, we performed flow cytometric analysis to assess differentiation efficiency. On day 6, definitive endoderm formation was confirmed by CXCR4 and CD117 across the three lines (Figure S3). By day 13, pancreatic progenitor specification was assessed through expression of NKX6-1, revealing 71.38% of cells were NKX6-1 positive (Figure S4). These analyses provided validation of efficient progression through definitive endoderm to pancreatic progenitor stages.

**Figure 1.**
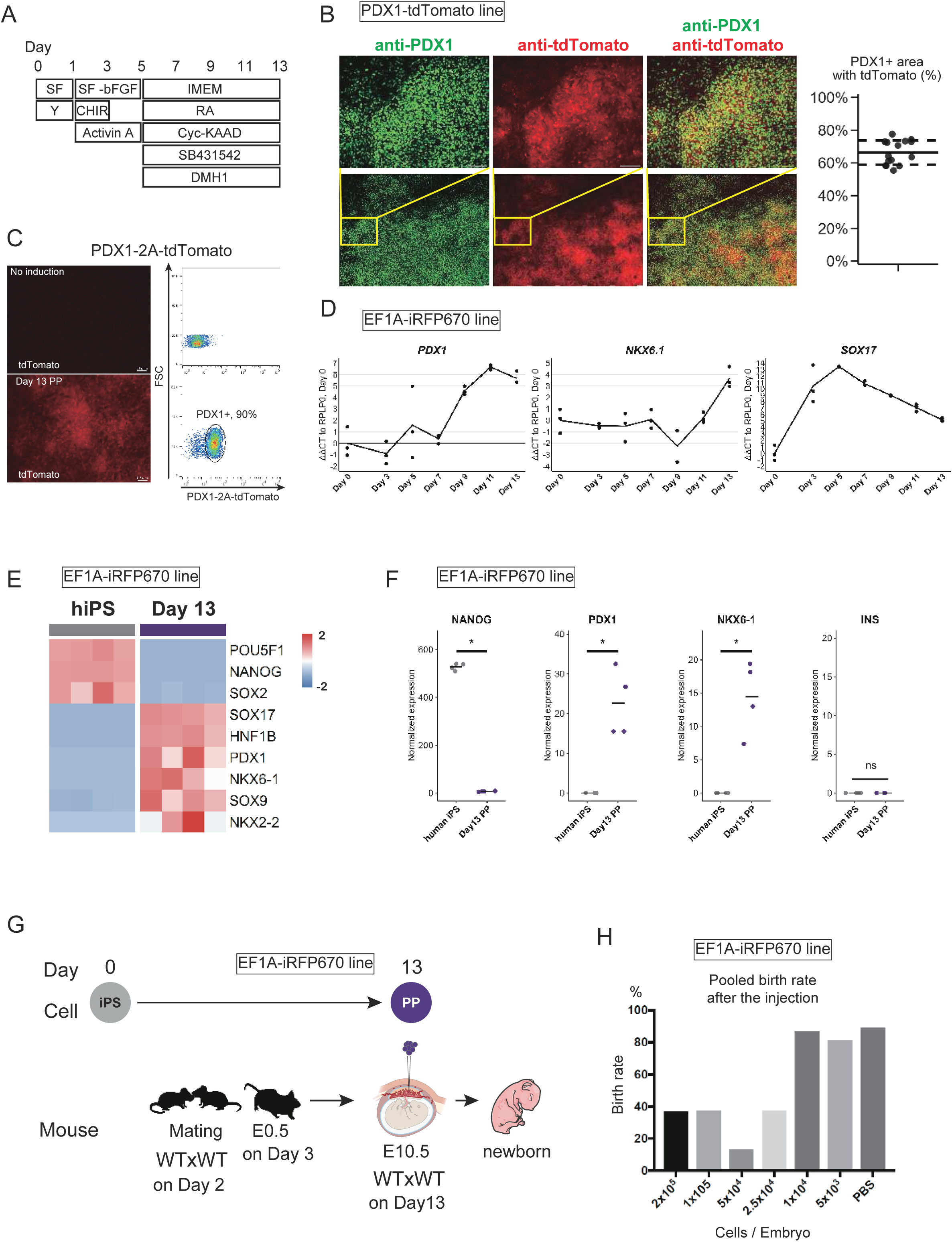
Differentiation of pancreatic progenitor and establishment of intraplacental injection. (A) Schematic representation of the stepwise differentiation protocol for generating PDX1+ pancreatic progenitors from human iPSCs. SF: StemFit medium, IMEM: Improved Minimum Essential Medium, CHIR: CHIR99021, Cyc-KAAD: Cyclopamine-KAAD, RA: retinoic acid. (B) Validation of PDX1-tdTomato reporter line. Representative immunofluorescence images showing co-localization of endogenous PDX1 protein (green) and tdTomato reporter (red) expression at day 13 of differentiation. Scale bar: 100 μm. Quantification shows 66.3% ± 7.4% co-localization efficiency after threshold. (C) Left: representative fluorescent microscopic image of differentiated PDX1-tdTomato line at day 13. Right: representative flow cytometric analysis of differentiated PDX1-tdTomato line at day 13. (D) Time-course expression analysis of pancreatic differentiation markers by RT-qPCR. Gene expression levels normalized to *RPLP0* and day 0 control. EEF1A1-iRFP670 line was used. Data shown as mean and each biological replicate, n = 3 independent experiments. (E) Transcriptomic comparison of iPSCs versus differentiated pancreatic progenitors at day 13. Heatmap showing differential expression of key developmental genes between hiPSCs (day 0) and day 13 pancreatic progenitors. Cell line: EEF1A1-iRFP670 line. Statistics: n = 4 biological replicates each condition. (F) Comparison of pancreatic differentiation markers’ normalized counts between undifferentiated hiPSCs and day 13 pancreatic progenitors. Cell line: EEF1A1-iRFP670 line. Statistics: n = 4, statistical analysis by DESeq2 Wald test, \**p* < 0.05, ns = not significant, data shown as individual points with mean. (G) Experimental workflow for intraplacental injection studies. Schematic showing cell differentiation timeline coordinated with mouse mating and embryonic injection at E9.5. Cell line: EEF1A1-iRFP670 line. (H) Optimization of cell injection dose for embryonic survival. Birth rates of WT embryos following intraplacental injection of human pancreatic progenitors at various cell concentrations. Cell line: EEF1A1-iRFP670 line. Sample sizes are detailed in Table S1.

For detailed transcriptomic profiling and subsequent injection experiments, we used the EEF1A1-iRFP670 line. Although iRFP670 expression intensity decreased approximately 10-fold after pancreatic differentiation (Figure S1E), the signal remained robust enough for cell tracking in subsequent transplantation experiments. RNA sequencing (RNA-seq) was performed to further characterize the differentiated cells. A total of 11,747 differentially expressed genes (DEGs; false discovery rate [FDR] < 0.01) were identified, and principal component analysis (PCA) showed that the gene expression profiles were completely separated before and after differentiation (Figure S5A). Genes associated with pancreatic development were upregulated in differentiated cells (Figure 1E and S5B), whereas the pluripotency markers *NANOG*, *SOX2*, and *POU5F1* were significantly downregulated (Figure 1E). Conversely, the pancreatic progenitor signature genes *PDX1*, *SOX9*, *HNF1B*, *NKX6-1*, *FOXA2*, and *HOXB4* were significantly upregulated (Figure 1E, 1F, and S5B). However, the pancreatic endocrine progenitor genes *NEUROD1* was not upregulated (Figure S5C).

Moreover, the pancreatic endocrine effector genes *INS*, *GCG*, *PPY*, and *SST* were not expressed (Figure 1F, Figure S5C). To investigate the differentiation potential of the pancreatic progenitors, we extended our culture protocol using a three-dimensional organoid system supplemented with betacellulin, exendin-4, and T3. After two additional weeks of culture, immunostaining revealed the presence of insulin-positive and glucagon-positive cells in organoids derived from all three cell lines (Figure S6), confirming that our PDX1-positive cells retained the capacity to further differentiate into pancreatic endocrine cells.

Taken together, these results demonstrated successful generation of PDX1-expressing pancreatic progenitor cells from multiple human iPSC lines with consistent efficiency and confirmed potential for endocrine differentiation. For subsequent injection experiments, we focused exclusively on the EEF1A1-iRFP670 line due to its superior cell tracking capabilities.

### Intraplacental injection of human PDX1-positive cells did not inhibit embryonic development at low concentrations

To set the cell number for injection, we first intraplacentally injected human pancreatic cells at various concentrations into wild-type (WT) embryos (Figure 1G, H, Table S1) at E10.5. EEF1A-iRFP670 cells were differentiated into pancreatic cells after 13 days of differentiation culture. On day 3, we mated the *Pdx1*^+/-^ mice to obtain E10.5 embryos on day 13 of culture (Figure 1G). The number of embryos was counted after injection. After injection, we waited for birth and counted the number of newborns to calculate the birth rate. The birth rate after injection was significantly lower (37, 37, 18, and 37%) at high concentrations (2 × 10^5^ to 2.5 × 10^4^ per embryo). On the other hand, low concentrations (1 × 10^4^ and 5 × 10^3^ per embryo) displayed birth rates comparable to that of the phosphate-buffered saline (PBS) injection control (>80%) (Figure 1H, Table S1). Frequent embryonic resorption was observed in mothers with low birth rates. Therefore, we set the concentration of cells at 1 × 10^4^ per embryo in all subsequent experiments.

### Human pancreatic cells extend the lifespan of *Pdx1*^-/-^ mice

Next, human pancreatic cells were intraplacentally injected into E9.5 or E10.5 embryos obtained from *Pdx1*^+/-^ mouse matings (Figure 2A). We confirmed that *Pdx1*^-/-^ mice were born according to the Mendelian rule (Table S2). Uninjected *Pdx1*^-/-^ newborns died a day after birth, as previously reported (Offield et al., 1996). On the other hand, human pancreatic cell-injected *Pdx1*^-/-^ newborns survived up to 10 days, and their lifespans were significantly longer than those of uninjected mice (*p* = 0.0013 in the log-rank test) (Figure 2B). Thus, *Pdx1*^-/-^ mice survived likely due to the contribution of human pancreatic cells. In our gross observations, the long-surviving *Pdx1*^-/-^ mice appeared small and thin compared with their WT and *Pdx1*^+/-^ littermates, indicating that the cells could not compensate for their growth abnormality (Figure 2C, D). In addition, pancreatic structures were not observed even in long-surviving *Pdx1*^-/-^ (Figure 2E).

**Figure 2.**
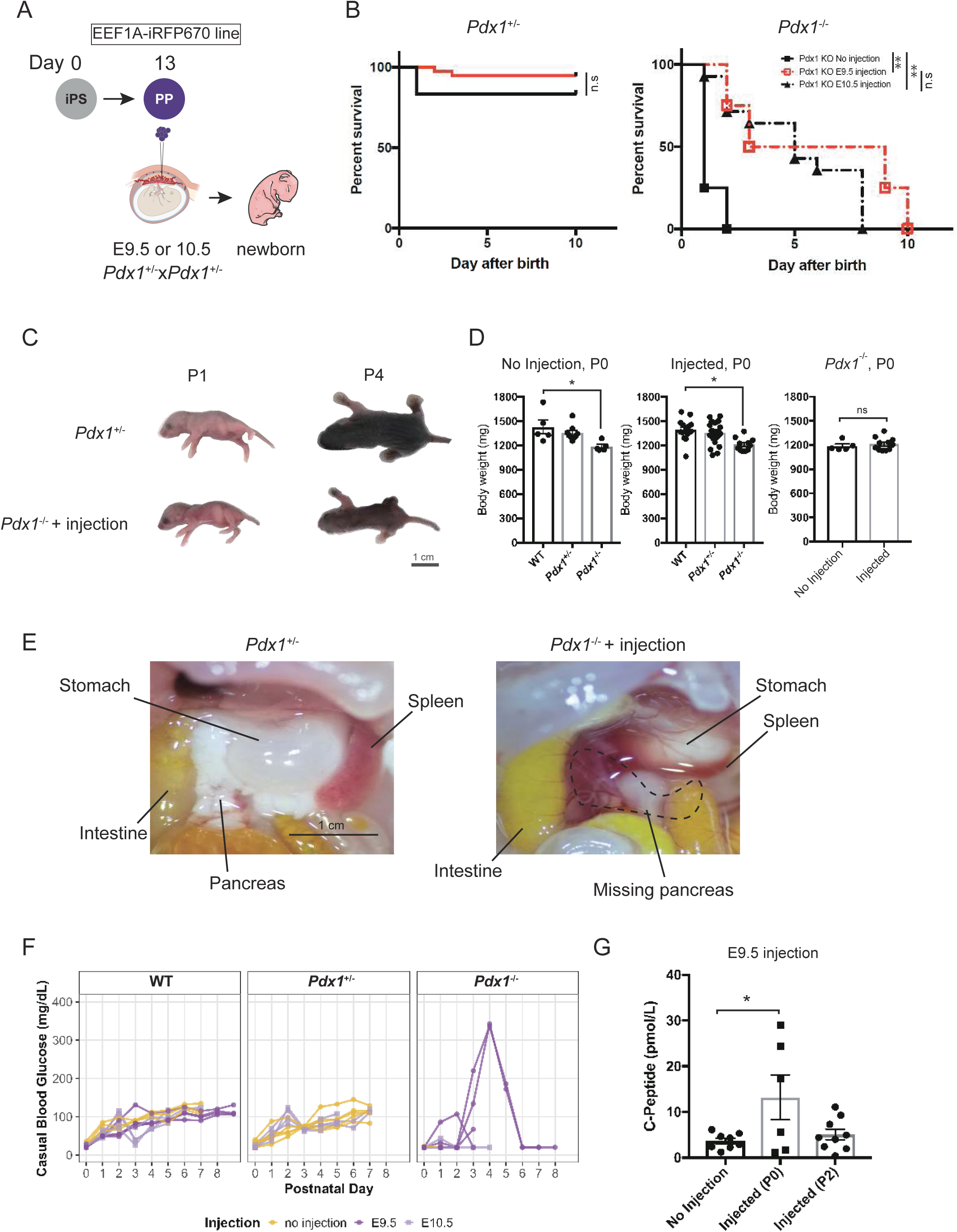
Injection of human pancreatic cells supports *Pdx1*^-/-^ newborn mice through insulin secretion. (A) Schematic showing intraplacental injection of human pancreatic progenitors into *Pdx1*^+/-^ mating embryos at E9.5 and E10.5. (B) Kaplan-Meier survival curves comparing *Pdx1*^+/-^ (left panel) and *Pdx1*^-/-^ mice (right panel) with and without human cell injection at E9.5 or E10.5. *Pdx1*^+/-^: no injection n = 10, injected n = 11; *Pdx1*^-/-^: no injection n =4, injected n = 11. Log-rank test, \**p* < 0.05, \*\**p* < 0.01, ns = not significant. (C) Representative photographs of *Pdx1*^+/-^ and *Pdx1*^-/-^ injected (E9.5 injection) mice at P1 and P4, showing growth differences. Scale bar = 1 cm. (D) Body weight comparison at P0. Data were shown as individual data points with mean ± SEM. Left and middle panel compare different genotypes. Right panel compares *Pdx1*^-/-^ mice with at E9.5 and without injection. No injection; WT, n = 5; *Pdx1*^+/-^, n = 8; *Pdx1*^-/-^, n = 5. Injected; WT, n = 15; *Pdx1*^+/-^, n = 25; *Pdx1*^-/-^, n = 11. Statistical analysis by one-way ANOVA with Tukey’s post-hoc test, \**p* < 0.05, \*\**p* < 0.01, ns = not significant. (E) Representative gross anatomy of internal organs at P0. Comparison between *Pdx1*^+/-^ (left) and *Pdx1*^-/-^ + injection at E9.5 (right) mice showing absence of pancreatic tissue in injected *Pdx1*^-/-^ mice. Scale bar = 1 cm. (F) Casual blood glucose measurements in no injection and injected mice (E9.5 injection and E10.5 injection). Each line represents an individual mouse. No injection; WT, n = 3; *Pdx1*^+/-^, n = 6; *Pdx1*^-/-^, n = 3. Injected at E10.5; WT, n = 4; *Pdx1*^+/-^, n = 4; *Pdx1*^-/-^, n = 3. Injected at E9.5; WT, n = 4; *Pdx1*^-/-^, n = 6. (G) Human C-peptide levels in serum of *Pdx1*^-/-^ mice with or without injection at E9.5. ELISA quantification comparing uninjected controls, injected mice at P0, and injected mice at P2. Data shown as individual points with mean ± SEM. Statistical analysis by one-way ANOVA with Tukey’s post-hoc test, \**p* < 0.05. No injection n = 8, Injected (P0) n = 6, and Injected (P2) n = 9. Experiments were done with EEF1A-iRFP670 line.

Therefore, although it did not correct the developmental abnormalities associated with their genotype, human pancreatic cell injection into *Pdx1*^-/-^ mice extended their lifespan.

### Human insulin production was detected in injected *Pdx1*^-/-^ mice

Given that *Pdx1*^-/-^ mice lack endogenous pancreatic β-cells and cannot produce insulin, we monitored blood glucose levels daily to assess potential functional human cell engraftment. Of the six *Pdx1*^-/-^ mice injected at E9.5, two mice showed increased blood glucose levels to 350 mg/dL on the fourth day of life, which decreased to below the detection limit (20 mg/dL) by the sixth day (Figure 2F). The other two did not show an increase in blood glucose and died on the third day of life. In contrast, mice injected at E10.5 showed no detectable glucose elevation throughout their survival period (Figure 2F). WT and *Pdx1*^+/-^ littermates maintained stable glucose levels around 100 mg/dL regardless of injection timing (Figure 2F). Based on these observations, subsequent experiments were performed using E9.5 injection timing.

To assess human insulin production, human C-peptide in *Pdx1*^-/-^ mouse serum was quantified using enzyme-linked immunosorbent assay (ELISA). Human C-peptide levels were significantly elevated at postnatal day 0 (P0) compared to uninjected controls (Figure 2G). By P2, while some individual animals retained detectable C-peptide levels, the group did not show statistically significant elevation compared to uninjected controls. The presence of human C-peptide suggests functional human β-cell activity in the injected mice.

These results demonstrated that injected human pancreatic progenitors can differentiate into insulin-producing cells in *Pdx1*^-/-^ mice.

### Human pancreatic cells engraft and express pancreatic genes in the duodenum of *Pdx1*^-/-^ newborns

Although the injected pancreatic cell-deficient mice survived significantly longer, pancreatic structures in their bodies were not observed (Figure 2E). Therefore, we investigated the ectopic engraftment of human pancreatic cells (Figure 3A). Several studies have reported that pancreatic cells can arise ectopically because of deficiencies in transcription factors essential for pancreatic development. Pancreatic endocrine cells have been observed in the spleen of *Ptf1a* knockout mice (Krapp et al., 1998). In addition, pancreatic cells have been found in the stomach, duodenum, and bile duct of *Hes1* knockout mice (Fukuda et al., 2006). We employed droplet digital PCR (ddPCR) to quantify human cell distribution in injected mice. This method enables absolute DNA quantification through partitioning PCR products into nanoliter-sized droplets, demonstrated previously to be effective for chimerism quantification (Suchy et al., 2024, 2022). We established a sensitive human DNA detection system using probes targeting primate-specific (*Caspase7*) and conserved mammalian (*Zeb2*) sequences. The assay was validated using defined mixtures of human and mouse DNA, confirming reliable detection down to 1% human DNA content (Figure S8). Analysis of multiple organs from P0 *Pdx1*^-/-^ newborns revealed that the duodenum contained significantly higher proportions of human DNA (2.8 ± 1.2%, mean ± SD, n = 8, *p* < 0.01) compared to uninjected controls, while other organs including truncated pancreas, kidney, intestine, liver, lungs, and spleen showed negligible human DNA content (Figure 3B).

**Figure 3.**
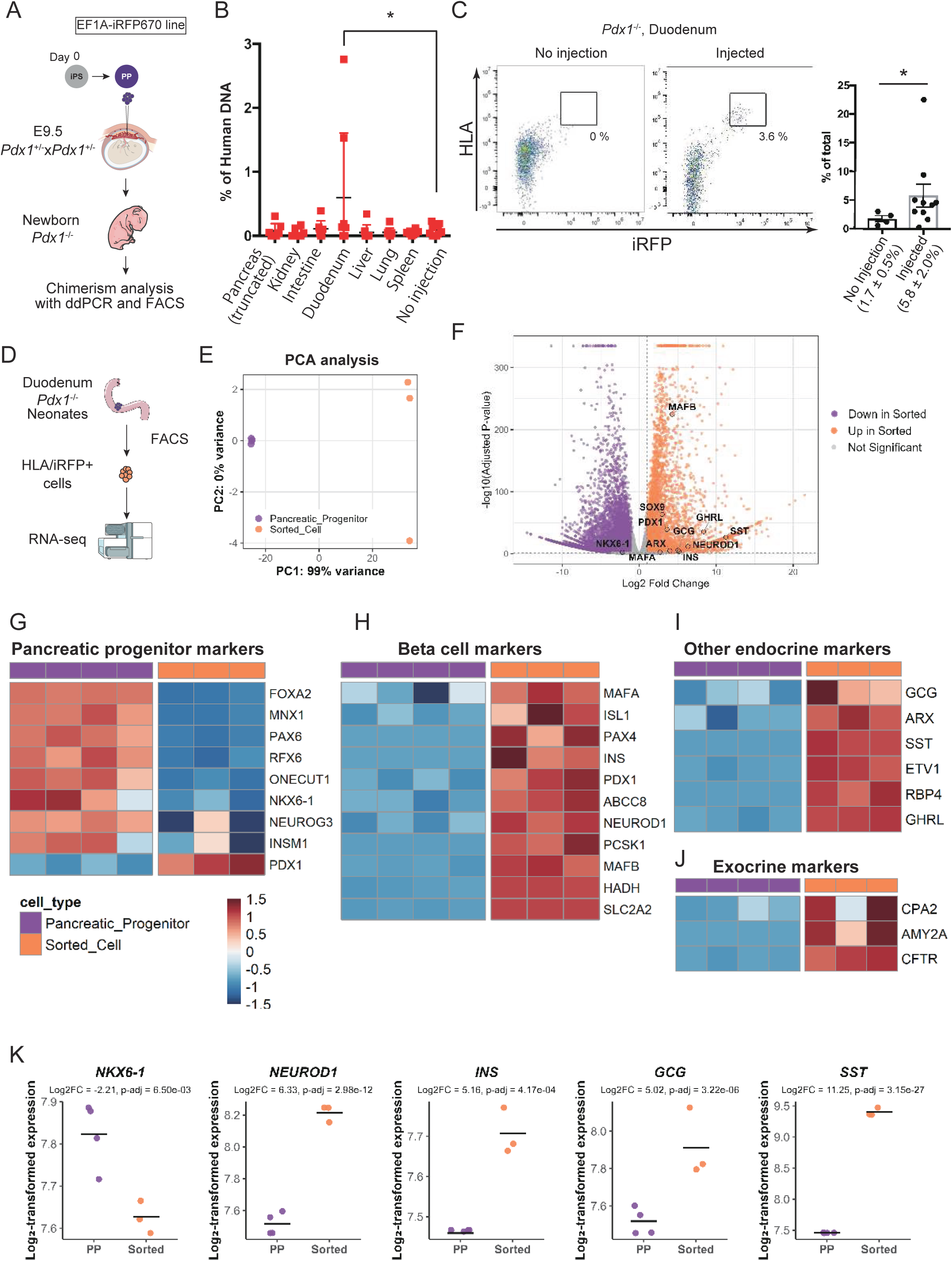
Human pancreatic cells engraft ectopically in the duodenum and undergo pancreatic differentiation. (A) Experimental workflow showing intraplacental injection at E9.5 followed by chimerism analysis using ddPCR and FACS at P0. (B) Quantification of human DNA distribution in various tissue by ddPCR. Percentage of human DNA detected in different tissues from injected *Pdx1*^-/-^ P0 newborns. n = 8 for all tissue, data shown as individual points with mean ± SEM. Statistical analysis by one-way ANOVA with Tukey’s post-hoc test, \**p* < 0.05. (C) Flow cytometric detection of human cells in duodenum of *Pdx1*^-/-^at P0. Representative FACS plots showing iRFP/HLA double-positive populations in uninjected versus injected *Pdx1*^-/-^ duodenum samples. Quantification shows the percentage of iRFP/HLA double-positive cells per DAPI-negative (live) cells. Data are shown as individual sample and mean ± SEM. No injection, n = 5, injected, n = 10. Statistical analysis by Welch’s t-test, \**p* < 0.05. (D) Experimental workflow showing RNA-seq analysis from sorted HLA/iRFP+ human cells. (E) PCA of transcriptomes from pancreatic progenitor cells (PP; purple, n = 4) and sorted engrafted cells (Sorted; orange, n = 3) showing clear separation. (F) Volcano plot of expressed genes between Sorted and PP populations. Upregulated (orange) and downregulated (purple) genes are indicated (|log2 fold change| > 1, adjusted *p* < 0.05; DESeq2 Wald test). (G) Heatmap showing pancreatic progenitor markers. (H) Heatmap showing upregulation of β-cell transcription factors and functional genes in Sorted cells. (I) Heatmap of α-cell (GCG, ARX), ε-cell (GHRL) and δ-cell (SST) marker expressions. (J) Heatmap of pancreatic exocrine marker expression. (K) Dot plots comparing VST-normalized expression levels for key pancreatic genes between PP and Sorted cells. Data are individual points with mean; n = 4 (PP) and n = 3 (Sorted); DESeq2 Wald test, p-values indicated. Experiments were done with EEF1A-iRFP670 line.

Flow cytometric analysis further confirmed human cell engraftment in the duodenum. We identified CD45.2 and CD31 double-negative (non-hematopoietic, non-endothelial) cells that were positive for both the transgenic human cell marker iRFP and human leukocyte antigen (HLA) in injected samples (Figure 3C). These human-specific marker-positive cells comprised 5.84 ± 2.00% of viable cells in the injected duodenum, significantly higher than in uninjected controls (1.78 ± 0.54%, *p* < 0.01).

Together, these ddPCR and flow cytometric analyses provided complementary evidence for specific human cell engraftment in the duodenum of injected *Pdx1*^-/-^ mice, with minimal distribution to other organs.

### Transcriptome analysis demonstrated human cells underwent differentiation after injection

To determine whether the engrafted human cells underwent functional maturation, we performed fluorescence-activated cell sorting (FACS) to isolate HLA and iRFP double-positive cells from the duodenum of injected *Pdx1*^-/-^ mice and compared their transcriptional profiles to the pancreatic progenitor cells used for injection donor (Figure 3D). RNA sequencing analysis was performed with reads mapped to the human genome. PCA demonstrated clear transcriptional divergence between the sorted cells and the progenitors, indicating substantial gene expression changes following engraftment (Figure 3E). DEG analysis revealed significant upregulation of pancreatic endocrine lineage genes in the engrafted cells compared to the starting population (Figure 3F). Engrafted human cells exhibited marked downregulation of pancreatic progenitor-associated genes, indicating progression beyond the progenitor stage (Figure 3G, K). Additionally, key β-cell transcription factors (*PDX1*, *NEUROD1*, *MAFA*, *MAFB*) and functional genes essential for glucose sensing (*GCK*), insulin biosynthesis (*INS*), insulin processing (*PCSK1*), and secretion machinery (*SLC30A8*, *ABCC8*, *KCNJ11*) were robustly upregulated, consistent with β-cell specification and functional maturation (Figure 3H, K). In addition to β-cell markers, elevated expression of α-cell (*GCG*, *ARX*), δ-cell (*SST*), and ε-cell (*GHRL*) genes was observed, whereas *PPY* transcripts were not detected (Figure 3I, K). Expression of acinar (*CPA2*, *AMY2A*) and ductal (*CFTR*) markers in the sorted cells further suggested the presence of a heterogeneous pancreatic lineage composition within the engrafted population (Figure 3J, K).

These transcriptomic analyses demonstrated that the human pancreatic cells engrafted in the duodenum maintained pancreatic identity and underwent differentiation toward more mature endocrine phenotypes including β-like cells, consistent with their ability to produce insulin and support extended survival of *Pdx1*^-/-^ mice.

### Transcriptome analysis reveals potential immune modulation in host duodenum after human cell engraftment

To investigate potential effects of human pancreatic progenitor cell engraftment on host tissue, we performed bulk RNA sequencing of duodenal tissue collected at P0 from *Pdx1^-/-^*mice with or without intraplacental human pancreatic cell injection. Reads were mapped to the mouse genome to exclude human cell-derived transcripts.

DEG analysis revealed substantial transcriptional alterations in the host duodenum following injection. In total, 1695 genes were significantly upregulated and 788 genes were downregulated in the injected group compared to uninjected controls (adjusted *p* < 0.05, |log2 fold change| > 1) (Figure S7A).

KEGG pathway enrichment analyses revealed that pathways related to innate immune responses such as neutrophil extracellular trap formation and cornified envelope were significantly upregulated. Conversely, gene sets associated with allograft rejection, autoimmune thyroid disease were significantly downregulated (Figure S7B). Representative heatmaps demonstrated coordinated upregulation of *Fpr1*, Complement *C3*, Histone component genes, suggesting an activated innate immune system in the duodenal tissue (Figure S7C). On the other hand, allograft rejection through T cells was suggested to be suppressed by coordinated downregulation of T cell receptor *Cd28*, *Fas*, and a series of MHC class I genes (Figure S7D). Interestingly, fat digestion and absorption were significantly downregulated (Figure S7B). Representative heatmaps demonstrated coordinated downregulation of intestinal absorption–related genes such as *Abcg8*, *Npc1l1*, *Apoa4*, and *Dgat1*, suggesting the incorporation of human cell might have impaired digestive function (Figure S7E).

Taken together, these transcriptomic findings suggest that xenogeneic cell injection might have altered the local epithelial and immune environment of the developing mouse intestine.

### IHC detected insulin-positive cell ectopic engraftment in *Pdx1*^-/-^ newborns duodenum

To directly visualize the engrafted human cells and their functional properties, we performed immunohistochemistry on duodenal sections from injected *Pdx1*^-/-^ newborns. Multiple insulin-positive cells were detected within the duodenal villi of injected samples across different individuals (Figure 4B, D). Since *Pdx1*^-/-^ mice lack endogenous pancreatic tissue and cannot produce insulin on their own, the presence of insulin-positive cells in the duodenum provided strong evidence that the injected human pancreatic progenitor cells not only engrafted but also differentiated into insulin-producing cells. Control duodenal sections from uninjected *Pdx1*^-/-^ mice showed no insulin-positive cells (Figure 4C, D).

**Figure 4.**
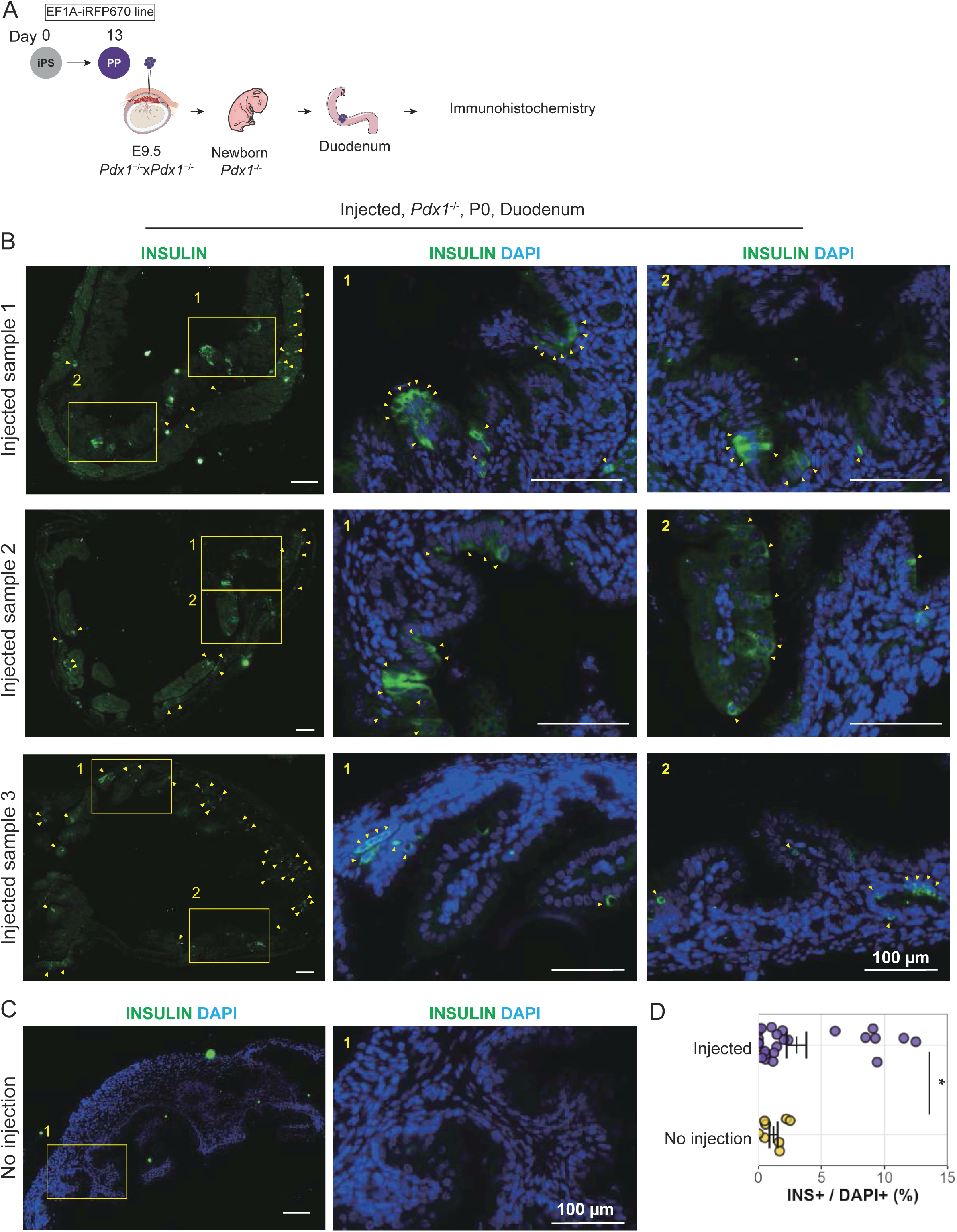
Immunohistochemical confirmation of insulin-positive human cells in the duodenum. (A) Experimental workflow for immunohistochemical analysis. Schematic showing intraplacental injection of human pancreatic progenitors into *Pdx1*^+/-^ × *Pdx1*^+/-^ matings at E9.5, followed by collection of duodenal tissue from *Pdx1*^-/-^ newborns at P0 for immunohistochemical analysis. (B) Representative immunofluorescence images of duodenal sections from three independent injected *Pdx1*^-/-^ newborns at P0. Insulin (green) with DAPI (blue) showing insulin-positive cells integrated within duodenal villi. Left panels show low magnification overview with boxed regions, right panels show higher magnification of numbered regions (1, 2) containing insulin-expressing cells. Scale bar = 100 μm. (C) Control duodenal section from an uninjected *Pdx1*^-/-^ newborn showing no detectable insulin-positive cells. Layout as in (B). (D) Quantification of insulin-positive cells as a percentage of total DAPI-positive cells per field. Eight fields were analyzed per sample. Data are shown as individual values with mean; Mann–Whitney U test, \**p* < 0.05. Experiments were done with EEF1A-iRFP670 line.

To rule out the possibility that human cells remained in the placenta and contributed to circulating human insulin, we analyzed placental tissues from injected embryos. WT mice were subjected to intraplacental injection and placental samples were collected at E18.5 via cesarean section to prevent maternal consumption. ddPCR analysis did not detect human DNA above background levels in placental samples (n = 13), confirming that injected cells did not significantly remain in the placenta (Figure S9B). RT-qPCR analysis of these placental samples showed no detectable expression of human *INS* (Figure S9C). The absence of insulin expression in placenta indicated that the human C-peptide detected in P0 serum samples must originate from human cells that were engrafted outside the placenta, most likely the insulin-positive cells we identified in the duodenum.

These immunohistochemical observations, together with our ddPCR, flow cytometry, and transcriptomic analysis, provided evidence that human pancreatic progenitor cells transplanted via intraplacental injection can engraft in the *Pdx1*^-/-^ mouse duodenum and differentiate into functional insulin-producing cells, which contributed to the extended survival of these pancreas-deficient mice.

## Discussion

Our study demonstrated the successful transplantation of human iPSC-derived pancreatic progenitor cells into mouse embryos via intraplacental injection. We established reproducible protocols for generating PDX1+ pancreatic progenitors with consistent differentiation efficiency and demonstrated their capacity to further differentiate into hormone-producing cells *in vitro*. Upon injection, the injected human cells predominantly localized to the duodenum where they expressed pancreatic genes and produced insulin, extending the lifespan of *Pdx1*^-/-^ mice from less than 1 day to up to 10 days after birth. Through multiple complementary analyses, including ddPCR, flow cytometry, immunohistochemistry, and transcriptome profiling, we confirmed the engraftment and functionality of human cells in the mouse neonatal environment. This *in vivo* functional validation provided evidence that specified human pancreatic progenitors can integrate into a developing mouse embryo and contribute to physiological functions, demonstrating the potential for creating interspecies chimeras with functional human-derived tissues.

Our approach differs from previous interspecies complementation studies that used blastocyst injection of pluripotent stem cells. Kobayashi et al. demonstrated that rat pluripotent stem cells injected into Pdx1-deficient mouse blastocysts generated structurally complete rat-derived pancreas in the host mouse, enabling survival beyond the neonatal period (Kobayashi et al., 2010). Similarly, Wu et al. showed efficient rat-mouse chimerism through blastocyst complementation, with chimeric contribution to multiple tissues (Wu et al., 2017). These studies utilized early-stage embryo injection of pluripotent cells, allowing broader developmental integration throughout embryogenesis. In contrast, our study introduced lineage-committed progenitor cells at a mid-gestation timepoint (E9.5), when the developmental trajectory is already substantially established. The use of lineage-committed progenitors rather than pluripotent stem cells offers several advantages. First, it minimizes the potential for unintended differentiation into other lineages, particularly germ cells and central nervous system, thus reducing ethical concerns associated with human-animal chimeras. Second, it provides a more directed approach for studying the integration and function of specific cell types. However, this strategy has limitations, as evidenced by the incomplete rescue of the *Pdx1*^-/-^ phenotype and lack of structural pancreatic organization. The contrasting outcomes between our approach and blastocyst complementation highlight the importance of developmental timing and cellular potency in determining the extent of functional integration in interspecies chimeras.

We observed the engraftment of human pancreatic progenitors in the duodenum. The duodenum expresses *Pdx1* during development (Offield et al., 1996), potentially providing a compatible developmental environment for PDX1+ human progenitor cells. This selective tropism is reminiscent of observations in *Hes1* knockout mice, where pancreatic cells develop ectopically in the stomach, duodenum, and bile duct (Fukuda et al., 2006), suggesting that epithelial cells in the digestive tract may share a developmental trajectory with pancreatic cells. The vascular connection between the placenta and developing organs likely plays a crucial role in cell distribution after intraplacental injection. At E9.5, the embryonic vascular system is established but still developing. The duodenum receives substantial blood flow through the superior mesenteric artery, which develops through extensive remodeling of the vitelline artery system and achieves functional capacity by this developmental stage (Downs et al., 1998). The omphalomesenteric circulation dominates over umbilical circulation until E9.5, directing substantial perfusion volume toward developing digestive organs (Hatch and Mukouyama, 2015; Walls et al., 2008). Our previous publication demonstrated that intraplacental injection of AAV9 resulted in widespread transduction across multiple organs, with particularly high efficiency in intestinal tissue, suggesting that the duodenum may indeed have advantageous blood flow characteristics (Gogoleva et al., 2023). This vascular architecture may facilitate the delivery of injected cells to the duodenal region. Further research using lineage tracing and real-time imaging could elucidate the precise mechanisms governing this selective cell engraftment.

Despite extending the lifespan of *Pdx1*^-/-^ mice, the transplanted human cells provided only partial functional rescue. Several factors likely contribute to this limited efficacy. First, the relatively low percentage of engrafted human cells is likely insufficient to fully compensate for the absence of an entire pancreas. Second, the engrafted cells did not organize into a structured pancreatic tissue with proper islet architecture, which is crucial for coordinated hormone secretion and glucose homeostasis. Immune-mediated rejection may also limit the long-term function of human cells. Our transcriptomic analysis of mouse duodenum suggested upregulation of genes associated with innate immune responses. These transcriptional changes suggest that xenogeneic cell presence may elicit a localized innate immune reaction, even in the absence of adaptive immune activation. Previous works have highlighted the importance of the CD47-SIRPα axis in xenograft rejection by macrophages (Azimzadeh et al., 2015; Ide et al., 2007), and innate immune mechanisms can act even in prenatal period. The use of SIRPα-humanized mice, which enable cross-species recognition of CD47, has been shown to reduce macrophage-mediated phagocytosis and enhance human cell engraftment *in vivo* (Jinnouchi et al., 2020). The absence of this compatibility in our model may contribute to the challenges in achieving sustained metabolic function, as evidenced by the variable glucose regulation patterns, human C-peptide levels and limited survival extension. Incorporating such immunomodulatory strategies will be critical for future efforts aimed at achieving stable and functional human cell integration.

Our study provided a proof-of-concept for human-animal chimera research using lineage-committed progenitor cells and intraplacental injection. This approach represents a technically accessible and ethically sound alternative to blastocyst complementation for studying human cell development and function *in vivo*. The high survival rate of injected embryos (>80% at optimal cell concentrations) overcomes previous limitations of *in utero* injection systems, enabling efficient delivery of human cells to developing embryos.

Several strategies could enhance the efficacy of this approach in future studies. Using immunodeficient recipients could reduce rejection and promote longer-term survival of human cells. Alternative recipient models with more selective defects, such as *Ngn3*^-/-^ mice lacking only endocrine pancreatic cells (Gradwohl et al., 2000), might allow more focused study of endocrine function without complete pancreatic agenesis. Additionally, engineering donor cells to express factors that enhance survival, such as anti-apoptotic genes or immunomodulatory molecules, could improve engraftment efficiency.

In conclusion, our study demonstrated the feasibility of generating interspecies chimeras with functional human pancreatic cells through *in utero* transplantation of lineage-committed progenitors. While the rescue of the *Pdx1*^-/-^ phenotype was incomplete, this work establishes a foundation for developing more sophisticated human-animal chimera models for studying human development and testing regenerative therapies. Future refinements in cell delivery, recipient selection, and donor cell engineering will likely enhance the utility of this approach for both basic research and translational applications.

## Experimental Procedures

### Animals

WT B6D2F1 (BDF1) mice were obtained from The Jackson Laboratory, Yokohama, Japan. Pdx1-LacZ heterozygous (*Pdx1*^+/-^) mice (Offield et al., 1996) were crossed with BDF1 mice for more than 10 generations to generate hybrids, as the better proliferation of hybrid strains was preferable. All mice were maintained under specific pathogen-free conditions at the University of Tsukuba Laboratory Animal Resource Center. All experiments were performed under relevant Japanese and institutional laws and guidelines and were approved by the University of Tsukuba Animal Ethics Committee (authorization number 23–039).

### Genotyping

Genotypes were assessed by PCR analysis of tail DNA. For genotyping, the primers for *Pdx1* loci were as follows: Pdx1_FW 5′-ATTGAGATGAGAACCGGCATG-3′, Pdx1-LacZ_FW 5′-TGTGAGCGAGTAACAACC-3′, and Pdx1_RV 5′-TTCATGCGACGGTTTTGGAAC-3′.

*Pdx1*^-/-^ embryos, the injection recipients, were obtained by mating *Pdx1*^+/-^ mice. The presence of a vaginal plug was checked the morning following mating, and the mice were considered to be carrying E0.5 embryos.

### Generation of PDX1-tdTomato and EEF1A1-iRFP670 iPS lines using genome editing technology

The PDX1-tdTomato and EEF1A1-iRFP670 iPS lines were generated using genome editing technology modified from our method published in a previous study (Nakade et al., 2023). Briefly, the expression vector for Cas9 nuclease and the guide RNA for cleaving the PDX1 gene (RDB20280: pX330-PDX1) were constructed by inserting annealed synthetic oligomers (5′-CACCgcagctcctgcctctcatcg-3′ and 5′-AAACcgatgagaggcaggagctgc-3′) into the Bbs I site of pX330-U6-Chimeric_BB-CBh-hSpCas9 vector (#42230; Addgene, Watertown, MA, USA; a gift from Prof. Feng Zhang). To construct the donor vector, a part of the human PDX1 gene corresponding to 1 kb up– and downstream of the stop codon was amplified by PCR using a primer set (PDX1_LAf: 5′-actgcaggttgcacccattcgcagg-3′ and PDX1_RAr: 5′-gccttgcaagatgttctcttcctcag-3′) and genomic DNA from WI38 cells as a template. The PCR product was cloned into the KpnI site of the pCRtk2×2NN vector (RDB18670) to generate pCRtk-PDX1 using an In-Fusion HD Cloning Kit (639648; Takara Bio Inc., Shiga, Japan). Subsequently, pCRtk2X2-PDX1 was amplified into two fragments by PCR using two primer sets (PDX1 LAr and p15AoriF and PDX1 RAf and p15AoriR). To prepare the reporter-selection marker, a part of the pUC-TEZ vector (RDB18672) harboring the P2A-tdTomato and EF1 alpha promoter-driven bleomycin resistance gene was amplified by PCR using the 2Af-1-16 and Bleo-R inf-puro primer set. PCR fragments were subjected to In-Fusion cloning to generate pUCtk-PDX1-TbsdEZ (RDB20281). The DNA sequences of all constructed plasmids were confirmed using a 3500 Genetic Analyzer (Applied Biosystems, Foster City, CA, USA).

To generate the EEF1A1-iRFP670 iPSC line, the expression vector for Cas9 nuclease and the guide RNA for cleaving the EEF1A1 gene, previously described as pX459-gEEF1A1 (RDB19570 deposited in the RIKEN DNA bank) (Nakade et al., 2023). The methods for constructing the donor vector for iRFP670 insertion were the same as those for PDX1-tdTomato, described previously. The iRFP670 sequence was cloned into pNLS-iRFP670 (#45466; Addgene) (Shcherbakova and Verkhusha, 2013). The resulting vector, pCRtk-EEF1A1-iRFP670-BSD (RDB20282), was used as the donor vector.

To generate knock-in cells, 5 µg of the donor vector and 1 µg of the Cas9-gRNA expression vector were introduced into 1 × 10^6^ cells of 1383D6 human iPSCs using an electroporator (NEPA21; Neppa Gene Co., Ltd., Chiba, Japan). Four days after electroporation, the electroporated cells underwent zeocin selection at 2 µg/mL. Since HSV-tk expression cassettes, designed outside left and right homology arms, were included in the donor vector, 30 µg/mL of ganciclovir was used to exclude clones with random insertions.

For the single-cell-derived colony cloning of PDX1-tdTomato clones, antibiotic-resistant cells were expanded to 100 cells/cm^2^ to isolate single-cell-derived colonies. Thirty-six isolated clones were examined for the presence of targeted insertion by PCR using PDX1_SeqUpLA: 5′-ATTTGCTGGCTCTCAGGTTGGGACT-3′ and PDX1_seqRA: 5′-GTTGAAGCCCCTCAGCCAGGAGCAG-3′. All 36 clones tested positive for the targeted insertion. For further experiments, we selected one clone as the PDX1-P2A-tdTomato line, which was deposited in the RIKEN Cell Bank as HPS4904. For the single-cell-derived colony cloning of the EEF1A1-iRFP670 line, antibiotic-resistant and iRFP670-positive cells were expanded to 100 cells/cm2 to isolate single-cell-derived colonies. Twelve Isolated clones were examined for the presence of targeted insertion by PCR using EEF1A1-up_FW: 5′-ACATGCTGGAGCCAAGTGCTA-3′ and EEF1A1-down_Rv: 5′-ATCACACAAACAGAAAGCATGTCC-3′. Eleven of the 12 clones were positive for targeted insertion. For further experiments, we selected one clone as the EEF1A1-iRFP670-TEZ line, which was deposited in the RIKEN Cell Bank as HPS5065.

While this knock-in reporter enables real-time visualization of PDX1 expression, the EF1α-promoter driven bleomycin resistance cassette incorporated into the construct may potentially influence endogenous PDX1 expression levels, localization, or function.

### Human iPSC culture

Undifferentiated human iPSCs were maintained in feeder-free StemFit AK02N medium (AK02N; Ajinomoto, Tokyo, Japan) to which iMatrix-511 (NP892-012; Nippi, Inc., Tokyo, Japan) and Y-27632 (Y0503; Sigma-Aldrich, St. Louis, MO, USA) was added 24 h after passage. All iPSCs were expanded to generate a master bank within the first five passages and served for the differentiation culture.

### Immunohistochemistry

For cultured cells, cells were fixed with 4% paraformaldehyde (FUJIFILM Wako Pure Chemical Corporation) for 10 minutes at room temperature. After washing with PBS, cells were permeabilized with 0.1% Triton X-100 (Wako) in PBS for 10 minutes and blocked with 1% bovine serum albumin (BSA; Wako) in PBS for 1 hour at room temperature. Primary antibodies were applied overnight at 4°C. Secondary antibodies conjugated with Alexa Fluor 488, 555, or 647 (Thermo Fisher Scientific) were applied for 1 hour at room temperature. Nuclei were counterstained with DAPI using Fluoro-KEEPER Antifade Reagent Non-Hardening Type (Nacalai Tesque). For tissue sections, samples were fixed in Mildform 10N (133-10311; FUJIFILM Wako Pure Chemical Corporation) and paraffinized. Paraffinized tissue blocks were sliced at a 4-μm thickness using a Sliding Microtome SM2000R (Leica Biosystems, Wetzlar, Germany). Samples were incubated in Antigen retrieval buffer (S1699; Agilent Technologies, Santa Clara, CA, USA) for 10 min at 110°C. After blocking with 5% goat serum/1% bovine serum albumin/PBS and Mouse on Mouse blocking reagent (Vector Laboratories, Newark, CA, USA), the samples were incubated with primary antibodies overnight at 4°C. Samples were washed in PBST for 10 minutes three times. Secondary antibodies conjugated with Alexa Fluor 488, 555, or 647 (Thermo Fisher Scientific) were applied for 1 hour at room temperature. Samples were washed in PBST for 10 minutes three times. Nuclei were counterstained with DAPI using Fluoro-KEEPER Antifade Reagent Non-Hardening Type (Nacalai Tesque).Tissue sections were mounted using VECTASHIELD Vibrance Antifade Mounting Medium (Vector Laboratories). Antibodies used were listed in Table S3. All samples were observed under a BIOREVO BZ-X800 microscope (Keyence, Osaka, Japan). All images were adjusted using Lightroom (Adobe, San Jose, CA, USA) and quantified using ImageJ software (NIH, Bethesda, MD, USA).

### Characterization of human iPSC lines

To validate pluripotency, undifferentiated human iPSCs were assessed for pluripotency marker expression through immunocytochemistry. The differentiation potential of established iPSC lines was evaluated through embryoid body formation. Approximately 1.2 × 10^6^ cells were suspended in StemFit AK02N supplemented with 12 μL of 10 mM Y-27632 and seeded into V-shaped 96-well plates to promote aggregate formation. After 2 days, cell aggregates were transferred to DMEM high glucose (Nacalai Tesque) supplemented with 10% fetal bovine serum (Biosera) and cultured for 8 days in suspension. Subsequently, embryoid bodies were plated onto 0.1% gelatin-coated dishes and maintained for an additional 8 days to allow spontaneous differentiation. On day 16, samples were fixed and processed for immunocytochemistry using lineage-specific markers representing the three germ layers. All antibodies used were listed in Table S3.

### Human iPSC differentiation culture to pancreatic progenitor

The culture medium used for the differentiation of PDX1-expressing cells comprised penicillin/streptomycin (P4333; Sigma-Aldrich) added to a basal medium (StemFit or Improved Minimum Essential Medium [IMEM]). On day 0, human iPSCs were seeded at 2,500 cells/cm^2^ in StemFit AK02N supplemented with iMatrix-411 (0.25 µg/cm^2^, NP892-041; Nippi, Inc.), iMatrix-511 (0.25 µg/cm^2^), and Y-27632 (10 µM). On day one, the medium was replaced by StemFit AK02N (A + B) supplemented with CHIR99021 (3 µM, #72052; STEMCELL Technologies, Vancouver, BC, Canada) and ActivinA (10 ng/mL) (725748-11-8; Fujifilm Wako Pure Chemical Corporation, Osaka, Japan). On day three, the medium was replaced by StemFit AK02N (A + B) supplemented with activin A (10 ng/mL). On day 5, the medium was changed to Modified IMEM with L-glutamine (A1048901; Thermo Fisher Scientific, Waltham, MA, USA) supplemented with B-27 (1% v/v, 17504044; Thermo Fisher Scientific), retinoic acid (2 µmol/L, R2625; Sigma-Aldrich), cyclopamine-KAAD (0.25 µmol/L, #20943; Cayman Chemical Company, Ann Arbor, MI, USA), SB431542 (10 µM, #72232; STEMCELL Technologies), and DMH-1 (10 µM, 4126/10; R&D Systems, Minneapolis, MN, USA).

For validation of PDX1 expression during pancreatic differentiation, cells at day 11 of differentiation were fixed and PDX1 protein was stained. Images were captured using a BZ-X800 fluorescence microscope (Keyence) with consistent exposure settings across samples at 20× magnification. Quantitative colocalization analysis between stained PDX1 and tdTomato reporter expression was performed using ImageJ software (version 1.54p, NIH, Bethesda, MD, USA) with the Coloc2 plugin. Twenty-five fields from differentiated cell cultures were analyzed. Colocalization was quantified using Manders threshold-corrected coefficients (tM1 and tM2) with Costes auto-threshold regression method to minimize background interference. Colocalization data was visualized using R software (version 4.4.2).

### Extended pancreatic organoid culture

To assess the endocrine differentiation potential of PDX1-positive progenitor cells, we employed a three-dimensional organoid culture system with maturation factors. On days 11-13 of the differentiation protocol, cells were harvested by washing with Dulbecco’s PBS followed by dissociation with 1 mL Tryple Express (Thermo Fisher Scientific, 2604-013) for 5 minutes at 37°C. Dissociated cells were collected, centrifuged at 200 × g for 3 minutes, and resuspended in pancreatic progenitor medium supplemented with 10 μM Y-27632. Cell suspensions were seeded into V-bottom 96-well plates (EZ-BindShut II, IWAKI, 4420-800LP) at 100 μL per well using multichannel pipettes, then centrifuged at 200 × g for 3 minutes to promote aggregate formation. On the following day, aggregates were cultured in pancreatic organoid medium consisting of Improved DMEM or Advanced DMEM supplemented with B-27 supplement (1% v/v), recombinant human betacellulin (20 ng/mL, 261-CE-010, R&D Systems), exendin-4 (2 μM, 055-09391, Wako), and 3,3’,5-triiodo-L-thyronine sodium salt (20 μM, 038-25541, Wako). Medium was changed every other day by aspirating half the volume and adding fresh organoid medium. After two weeks of organoid culture, samples were collected for histological analysis by embedding in IP Gell (Genostaff) followed by fixation with 4% paraformaldehyde.

### Flow cytometric analysis

For evaluation of pancreatic differentiation efficiency, cells were harvested at specific timepoints and processed for flow cytometric analysis. At day 3 of differentiation, definitive endoderm formation was assessed using antibodies against CD117 (c-Kit) and CXCR4 (CD184). Cells were dissociated using TrypLE Express (Thermo Fisher Scientific) for 15 minutes at 37°C, collected in PBS containing 10% FBS, and centrifuged at 200 × g for 3 minutes. Cell pellets were resuspended in FACS buffer (1% FBS in PBS) and incubated with PE anti-human CD117 (BioLegend) and APC anti-human CXCR4 (BioLegend) antibodies at 5 μL per million cells for 30 minutes at room temperature. Following two washes with FACS buffer, cells were resuspended in 200 μL FACS buffer containing DAPI (DOJINDO) for viability assessment.

For pancreatic progenitor evaluation at days 11-14, NKX6-1 expression was analyzed using intracellular staining. Cells were dissociated with Accutase (Nacalai Tesque) for 15 minutes at 37°C, fixed with BD Cytofix fixation buffer (Cat. No. 554655) for 15-30 minutes at 4°C, and permeabilized using BD Phosflow Perm Buffer III (Cat. No. 558050) for 30 minutes on ice. Cells were then incubated with Alexa Fluor 488 Mouse Anti-NKX6-1 antibody (Cat. No. 567608, BD Biosciences) for 15 minutes on ice, followed by two washes with FACS buffer before analysis.

For analysis of human cell engraftment in injected mice, the duodenum of newborn mice was dissociated using collagenase type V (032-17854; FUJIFILM Wako Pure Chemical Corporation) following a published protocol (Uniken Venema et al., 2022). Single-cell suspensions were filtered through 40-μm cell strainers and resuspended in PBS containing 2% fetal bovine serum. Human cells were identified by dual-positive expression of HLA and the constitutive iRFP670 reporter, with additional exclusion of hematopoietic (CD45.2+) and endothelial (CD31+) lineages to focus on non-vascular engrafted cells.

Data acquisition was performed using CytoFLEX (Beckman Coulter). Data analysis was conducted using CytoExpert (Beckman Coulter) and FlowJo software (TreeStar Inc., Ashland, OR, USA). All antibodies used are summarized in Table S3.

### Reverse transcription-quantitative PCR

Total RNA was collected using an ISOGEN (Cat. No. 311-02501; Nippon Gene, Tokyo, Japan). cDNA was synthesized using a QuantiTect Reverse Transcription Kit (Cat. No. 205313; Qiagen, Hilden, Germany). Gene expression levels were determined using RT-qPCR performed on a Thermal Cycler Dice Real Time System Single TP850 (Takara Bio Inc.) using THUNDERBIRD® SYBR® qPCR Mix (QPS-201; TOYOBO Co., Ltd., Tokyo, Japan). mRNA levels were normalized to those of RPLP0. The primer sequences used were summarized in Table S4.

### RNA sequence analysis of human iPSC differentiation

Total RNA was extracted using TRIzol (Thermo Fisher Scientific) following standard protocols. An RNA-seq library was prepared using an NEBNext Ultra II Directional RNA Library Prep Kit (New England Biolabs, Ipswich, MA, USA) after ribosomal RNA (rRNA) depletion using an NEBNext rRNA Depletion Kit (New England Biolabs). Paired-end (2 × 36 bases) sequencing was performed using a NextSeq500 platform (Illumina, San Diego, CA, USA). The FASTQ files were imported into the CLC Genomics Workbench (version 10.1.1; Qiagen), and the sequence reads were mapped to the human reference genome (hg38). Gene expression was calculated as the total read count normalized to transcripts per million. DEGs were extracted using the criterion of FDR-corrected p < 0.05.

### RNA sequence analysis of chimeric duodenum

Paired-end (2 × 36 bases) sequencing was performed using the same protocol for the iPSC differentiation experiment. FASTQ files were imported into STAR 2.7.11b. Sequence reads were mapped onto the mouse reference genome (mm10) without allowing a single mismatch (--outFilterMismatchNmax 0) or multimapping (--outFilterMultimapNmax 1). DEGs were extracted using DESeq2 in R (Love et al., 2014) using the criteria of *p* < 0.05 and log2 fold change > 2. GO analysis was performed using clusterprofiler (Wu et al., 2021). Clustering heatmaps were constructed using R software.

### FACS sorting and RNA sequencing of human cells

For transcriptome analysis of engrafted human cells, duodenal tissues from injected *Pdx1*^-/-^newborns were harvested at P0 and processed for FACS. Tissues were dissociated using collagenase type V (032-17854; FUJIFILM Wako Pure Chemical Corporation) as described above, followed by filtration through 40-μm cell strainers to obtain single-cell suspensions. Dead cells were excluded using DAPI staining, and human cells were identified by dual-positive expression of HLA and the iRFP670 reporter. Sorting was performed using a MoFlo XDP sorter (Beckman Coulter). Control samples from uninjected *Pdx1*^-/-^ mice were processed in parallel to account for potential background signals.

RNA sequencing libraries were prepared using the same protocol as above. Paired-end sequencing (2 × 150 bases) was performed on a NextSeq500 platform (Illumina, San Diego, CA, USA). Total RNA was extracted using ISOGEN-LS (NIPPON GENE). Sequence reads were processed using STAR aligner (version 2.7.11b) with Two-pass mapping mode. Reads were mapped exclusively to the human reference genome (hg38) using parameters (outFilterMismatchNoverReadLmax 0.06, outFilterMultimapNmax 10, outFilterMatchNminOverLread 0.3, outFilterScoreMinOverLread 0.3, alignSJoverhangMin 8, alignSJDBoverhangMin 3). DEG analysis was performed using DESeq2 (version 1.34.0) in R, with genes showing log2 fold change > 2 and adjusted p-value < 0.05 considered significantly differentially expressed. DEGs were listed in Table S5.

### Intraplacental injection

Intraplacental injection was performed with prior protocols (Jeon et al., 2021). Shortly before injection, the differentiated cells were dissociated using Accutase (12679-54; Nacalai Tesque, Kyoto, Japan) for 15 min at 37°C and washed in PBS twice. The cells were then suspended in cold PBS. Pregnant mice bearing E9.5 embryos were anesthetized with isoflurane while resting on a 37°C heating plate. The uterus was carefully exposed through an abdominal incision. The donor cell suspension was loaded into the injection needle through mouth pipetting using rubber tube (Drummond, 68-0449-50) and subsequently installed to a microinjector (FemtoJet; Eppendorf, Hamburg, Germany). Each placenta was punctured by an injection needle to a depth of 2 mm, and the donor cell suspension was injected at an injection pressure of 60 hPa. After injection, the uterus was carefully repositioned, and the incision was securely closed. After surgery, the mice were placed on a plastic bag containing warm water to prevent hypothermia.

### Survival analysis

Individual identification of newborn mice was established by tattooing their limbs on the day of birth. Genotyping was performed using tail-tip samples. Maternal care was enhanced by supplementing the maternal diet with CMF sprouts (Oriental Yeast; Tokyo, Japan). Newborn survival was checked daily until all *Pdx1*^-/-^ mice died. The Kaplan–Meier curve was constructed using GraphPad PRISM 6 (GraphPad Software, La Jolla, CA, USA).

### Glucose measurement

A small amount of blood was collected daily via facial vein puncture. Blood glucose levels were measured using a glucometer (Terumo, Tokyo, Japan). The minimum detection limit is 20 mg/dL and all glucose levels lower than the limit are recorded as 20 in our figure 2F.

### ELISA assay

ELISA was conducted using a Mercodia Ultrasensitive C-peptide ELISA kit (#10-1141-01; Mercodia, Uppsala, Sweden). Blood samples were collected from P0 and P2 *Pdx1*^-/-^ mice by decapitation. Approximately 25 μL of serum was obtained from each animal and diluted 4-fold with the kit’s calibrator diluent to achieve the required 100 μL sample volume for the assay. For the ELISA procedure, calibrator, control, and diluted sample volumes of 100 μL each were added to their respective wells according to the manufacturer’s protocol. The plate was incubated on a shaker for 1 h at 25°C and then washed six times with 350 μL of wash buffer. Subsequently, 200 μL of the enzyme conjugate solution was added to each well, followed by another 1 hour incubation on the plate shaker at room temperature. The plate was then rewashed six times, and 200 μL of TMB substrate was added to each well, followed by a 30 minutes incubation on the bench at room temperature protected from light. Next, 50 μL of stop solution was mixed into each well to terminate the reaction. Optical density readings were obtained at 450 nm using a microplate reader (SpectraMax M5, Molecular Devices) within 30 min. Baseline signals from uninjected control samples were used as assay controls, and only values significantly above this baseline (*p* < 0.05, Student’s t-test) were interpreted as positive detection of human C-peptide. All samples were measured in duplicate, and the mean values were used for analysis.

### ddPCR analysis

For ddPCR analysis, a QX200 Droplet Digital PCR system (Bio-Rad Laboratories, Hercules, CA, USA) was used. Briefly, to prepare DNA, tissue lysis was conducted in 10% proteinase K solution in a 1% SDS buffer. The samples were incubated at 55°C overnight. The following morning, another round of lysis buffer was added, and incubation was continued for a few additional hours. Proteinase K inactivation was accomplished at 75°C for 10 min, followed by extraction of 50 µL from each sample. To precipitate SDS, 5 µL of 1 M KCl was added, and the samples were briefly centrifuged at 3,000 × g for 1 min at 4°C. The reaction mixture was prepared using a custom ddPCR assay (#10031276 for FAM and #10031279 for HEX; Bio-Rad Laboratories) and ddPCR Supermix for Probes (no dUTP) (#186-3023; Bio-Rad Laboratories) according to the manufacturer’s instructions. Droplet formation was performed using droplet generator oil (#186-3005; Bio-Rad Laboratories). The subsequent PCR reaction involved enzyme activation at 95°C for 10 min, denaturation at 94°C for 30 s, and annealing/extension at 60°C for 1 min. This PCR cycle was repeated 50 times, and enzyme deactivation was achieved at 98°C for 10 min, with a final holding step at 12°C. QuantaSoft software (Bio-Rad Laboratories, version 1.7) was used to quantify the absolute number of DNA copies in all wells for data acquisition and analysis. Mouse DNA and human DNA were used to supervise thresholds.

The sequences of primers and probes used are as follows:

*chCasp7*:

Fw-GAAGAAGAAAAATGTCACCATGCG, Rv-TTTCAAAATTCATGTTGTACTGATATGTAG, Probe-AGACCACCCGGGACCGAGTGCCT

*Zeb2*:

Fw-GGATGGGGAATGCAGCTCTT, Rv-AGTGCGGCAGAATACAGCA, Probe-TGATGGGTTGTGAAGGCAGCTGCACCT

For validation of these primers and probes, human DNA from EEF1A-iRFP670 and mouse DNA from tail tips were used. DNA was mixed with known ratio (0, 1, 10, 50, 100% human DNA) and served to benchmark its accuracy (Figure S8).

### Placental tissue analysis

Pregnant WT mice that had received intraplacental injection of human pancreatic progenitor cells at E9.5 were subjected to cesarean section at embryonic day 18.5 to collect placental tissues before potential maternal consumption. Placentas were carefully dissected from the uterine wall, rinsed with cold PBS to remove maternal blood, and immediately processed for downstream analysis. For DNA extraction, placental tissues were minced and subjected to tissue lysis before proceeding to ddPCR analysis as described above.

For RNA analysis, total RNA was extracted using ISOGEN (NIPPON GENE) following the manufacturer’s protocol. Quantitative RT-PCR was performed to detect human insulin (INS) mRNA expression. Human INS ORF plasmid was served as positive control. Duodenum from no-injection newborn was served as negative control.

### Statistical analysis

Statistical analyses were performed using R (version 4.4.2) and GraphPad PRISM 6 software. The statistical significance of survival time was determined using the log-rank test. The statistical significance of the differences between three or more groups was determined using one-way ANOVA followed by Tukey–Kramer’s multiple comparisons. The statistical significance of the quantification of the immunohistochemistry was determined using Mann-Whitney U test. Statistical methods specific to RNA-sequencing data are described in their respective sections. The statistical significance of the FACS analysis was determined using Welch’s t-test. Statistical significance was considered at *p* < 0.05.

## Resource availability

### Lead contact

Michito Hamada, Ph.D., Department of Anatomy and Embryology, Faculty of Medicine, University of Tsukuba, 1-1-1 Tennodai, Tsukuba 305-8575, Japan; Phone: +81-298-53-7516, Fax: +81-298-53-6965

### Materials availability

This study did not generate new unique reagents.

### Data and code availability

RNAseq data was deposited as accession number DRR545160-DRR545181.

Additional metadata, culture details and basic characterization of the cell lines were deposited to RIKEN Cell Bank, as HPS4904 (PDX1-P2A-tdTomato) and HPS5065 (EEF1A1-iRFP670).

## Acknowledgments

We thank Dr. Yoshihiro Miwa, Dr. Koji Nakade, Dr. Kenichi Nakashima, and Mrs. Yasuko Hemmi for their technical support in generating PDX-P2A-tdTomato and EEF1A1-iRFP670 hiPSC lines. We thank Dr. Satoshi Yamazaki for providing *Pdx1*^LacZ/+^ mice and technical support for the flow cytometric analysis. We are grateful to all former and current members of the Department of Anatomy and Embryology for their support in all aspects. This work was supported by JSPS KAKENHI (Grant Number JP22H04922, 23K05586, 23K18208), JST (Grant Number JPMJPF2017), AMED (Grant number 23bm1423010h0001), and Research Fellowships of Japan Society for the Promotion of Science for Young Scientists (23KJ0238).

## Author contributions

Conceptualization, S.T., Y.H., M.H.; data curation, A.W.; formal analysis, A.W., Z.J.S.; funding acquisition, S.T., A.W., Y.H., H.N.; investigation, A.W., H.J., Z.J.S., T.H., C.W.L., N.G., A.N., Y.H., F.S. Y.A.; methodology, A.W., H.J., M.H., Y.H., F.S.; project administration, A.W., M.H., S.T.; resources, S.T., Y.H., H.N.; software, A.W., T.H.; supervision, S.T., M.H.; validation, A.W., H.J., Z.J.S., T.H., C.W.L., N.G., A.N., Y.H., F.S.; visualization, A.W., Z.J.S., M.H.; writing-original draft, A.W.; writing-review & editing, A.W., H.J., Z.J.S., T.H., C.W.L., N.G., A.N., F.S., H.N., Y.H., M.H., S.T.

## Declaration of generative AI and AI-assisted technologies in the writing process

During the preparation of this work, the authors used Claude 4 Sonnet in order to improve language and readability. After using this tool/service, the authors reviewed and edited the content as needed and take full responsibility for the content of the publication.

