## Supplemental data for "Intraplacental injection of human iPSC-derived PDX1+ pancreatic progenitors prolongs Pdx1-deficient mice survival"

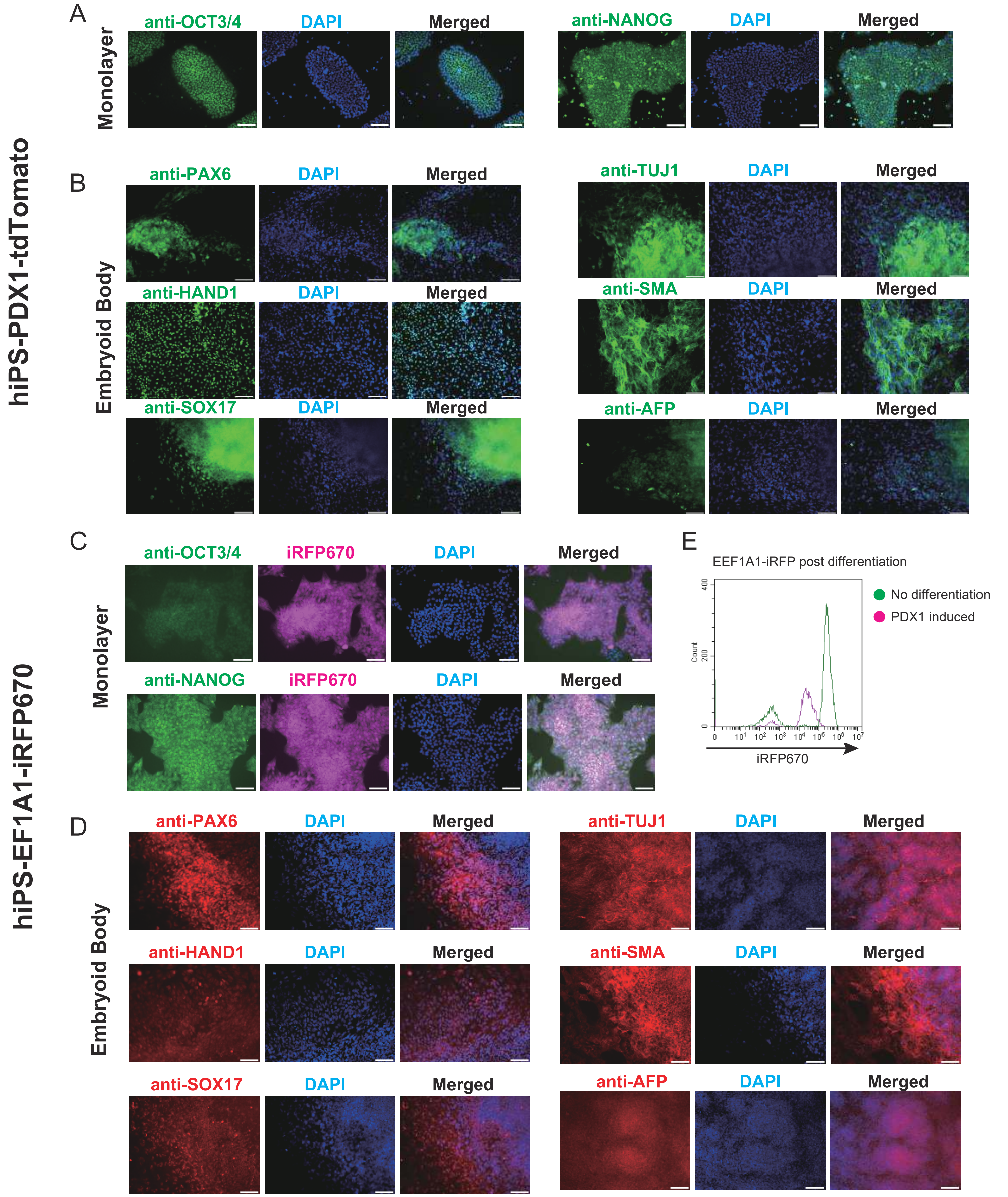


### Figure S1 Comprehensive characterization of newly established human iPSC lines

(A and C) Immunohistochemistry of undifferentiated PDX1-tdTomato and EEF1A1-iRFP670 lines showing robust expression of pluripotency markers OCT3/4 and NANOG. Scale bars: 100 μm. (B and D) Immunohistochemistry of embryoid body after 16 days of spontaneous differentiation to assess differentiation potential. Both lines expressed markers representing all three germ layers: ectoderm (PAX6: early neural specification, TUJ1: neuronal differentiation), mesoderm (HAND1: cardiac/extraembryonic mesoderm, SMA: smooth muscle), and endoderm (SOX17: definitive endoderm, AFP: hepatic specification). Scale bars: 100 μ m. (E) FCM histogram of EEF1A-iRFP670 with or without pancreatic progenitor differentiation.

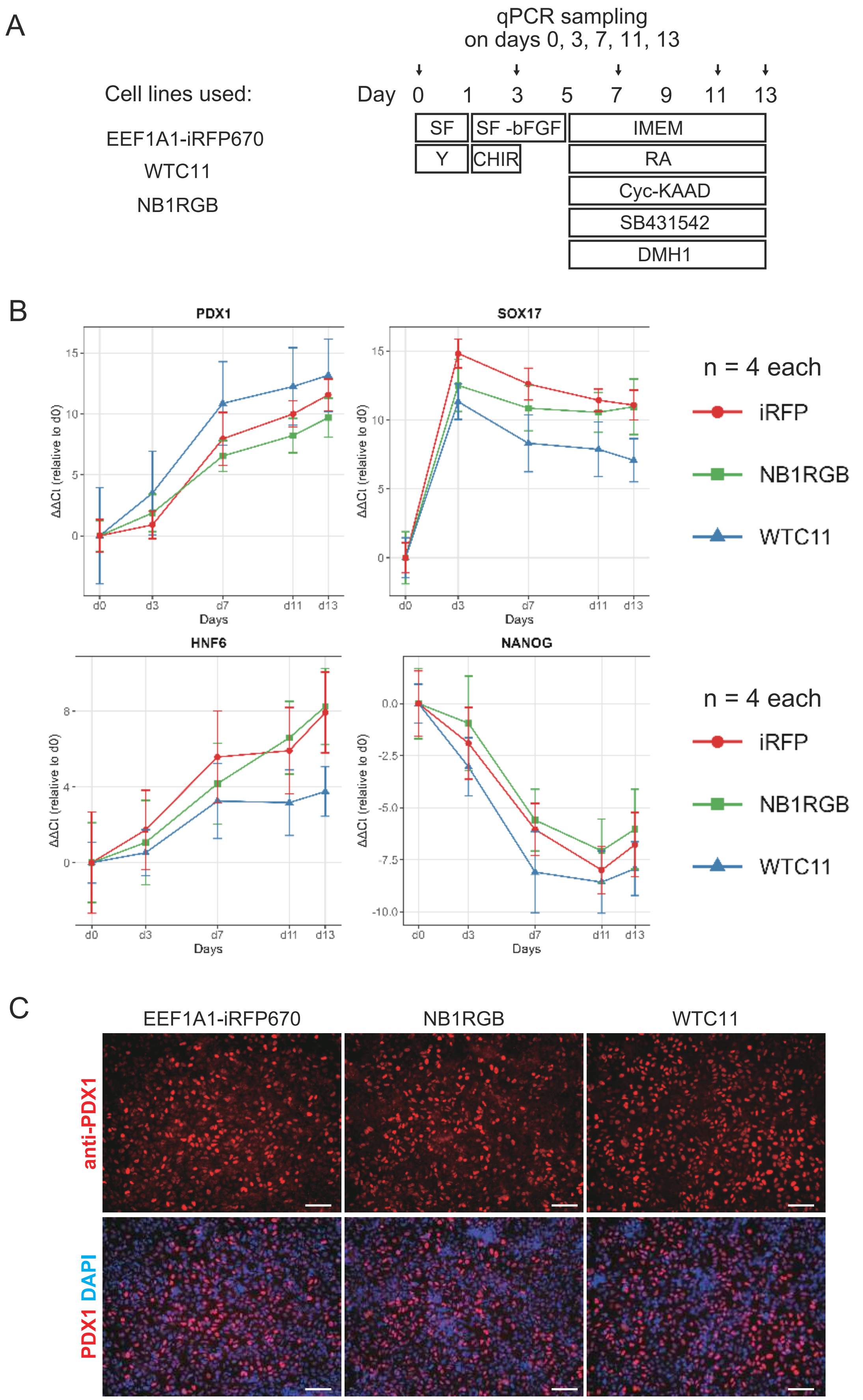


### Figure S2 Reproducible pancreatic differentiation across multiple iPSC lines

(A) Schema of the differentiation protocol and the RT-qPCR timecourse experiment for differentiating PDX1-expressing pancreatic progenitor cells from three different human iPSC lines. (B) Relative gene expression levels of *PDX1*, *SOX17*, *HNF6*, *NANOG* on day 0, day 3, day 7, day 11, day 13 of the differentiation. Data were normalized to *RPLP0* and day 0, and shown as mean +/- SE. (C) Immunostaining analysis of PDX1 expression of the three cell lines on day 13 of the differentiation. Scale bars: 100 µm.

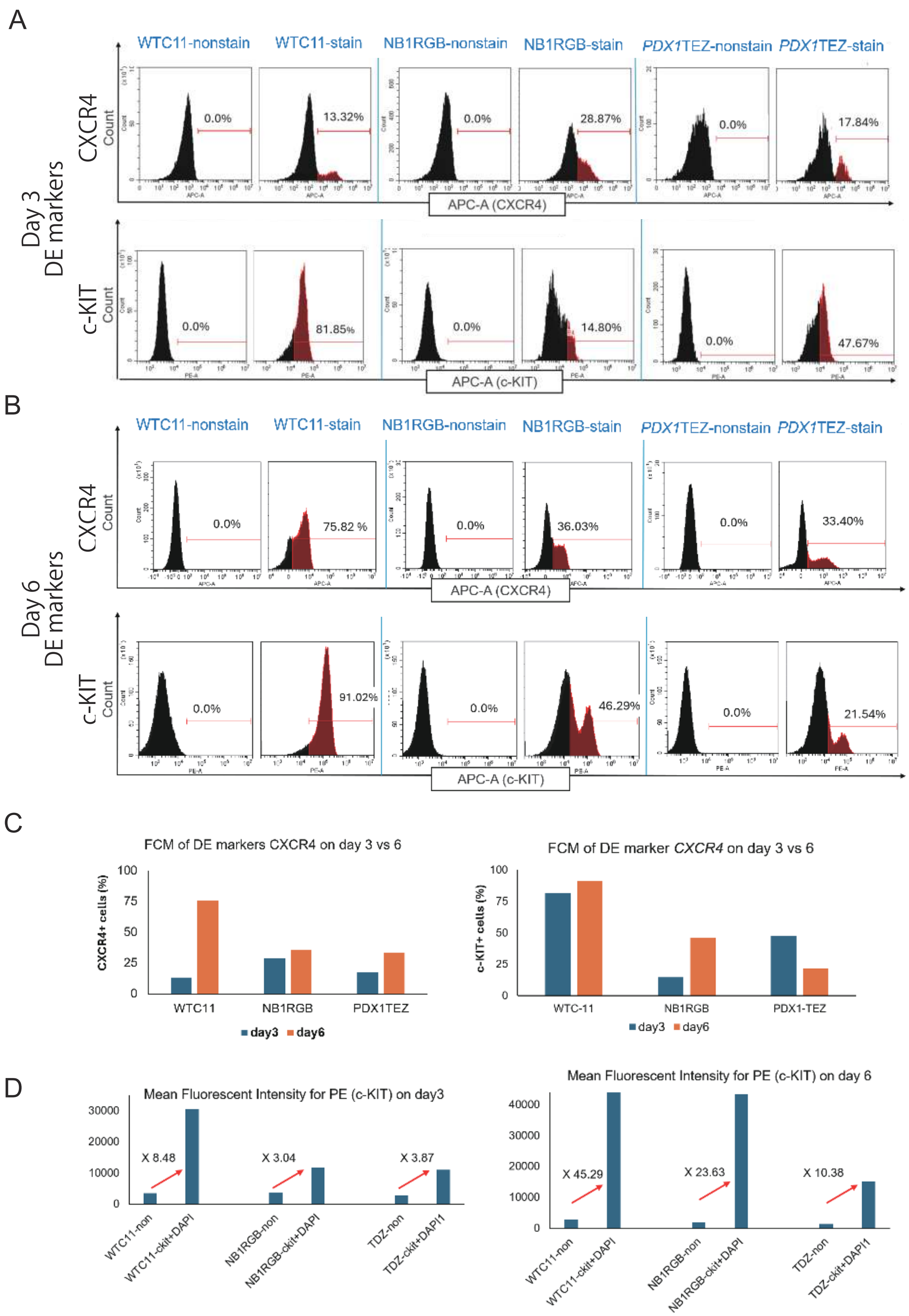


### Figure S3 Flow cytometric analysis of definitive endoderm (DE) markers during pancreatic differentiation of three iPSC lines

(A, B) Flow cytometric analysis of CXCR4 and c-KIT expression in WTC11, NB1RGB, and PDX1-tdTomato lines during DE differentiation. Representative histograms at day 3 (A) and day 6 (B). (C) Quantification of CXCR4+ cell percentages. (D) Mean fluorescence intensity of c-KIT expression at day 3 (left) and day 6 (right). The "non" bar indicates the non-stained control sample, which serves as the negative baseline for MFI measurement.

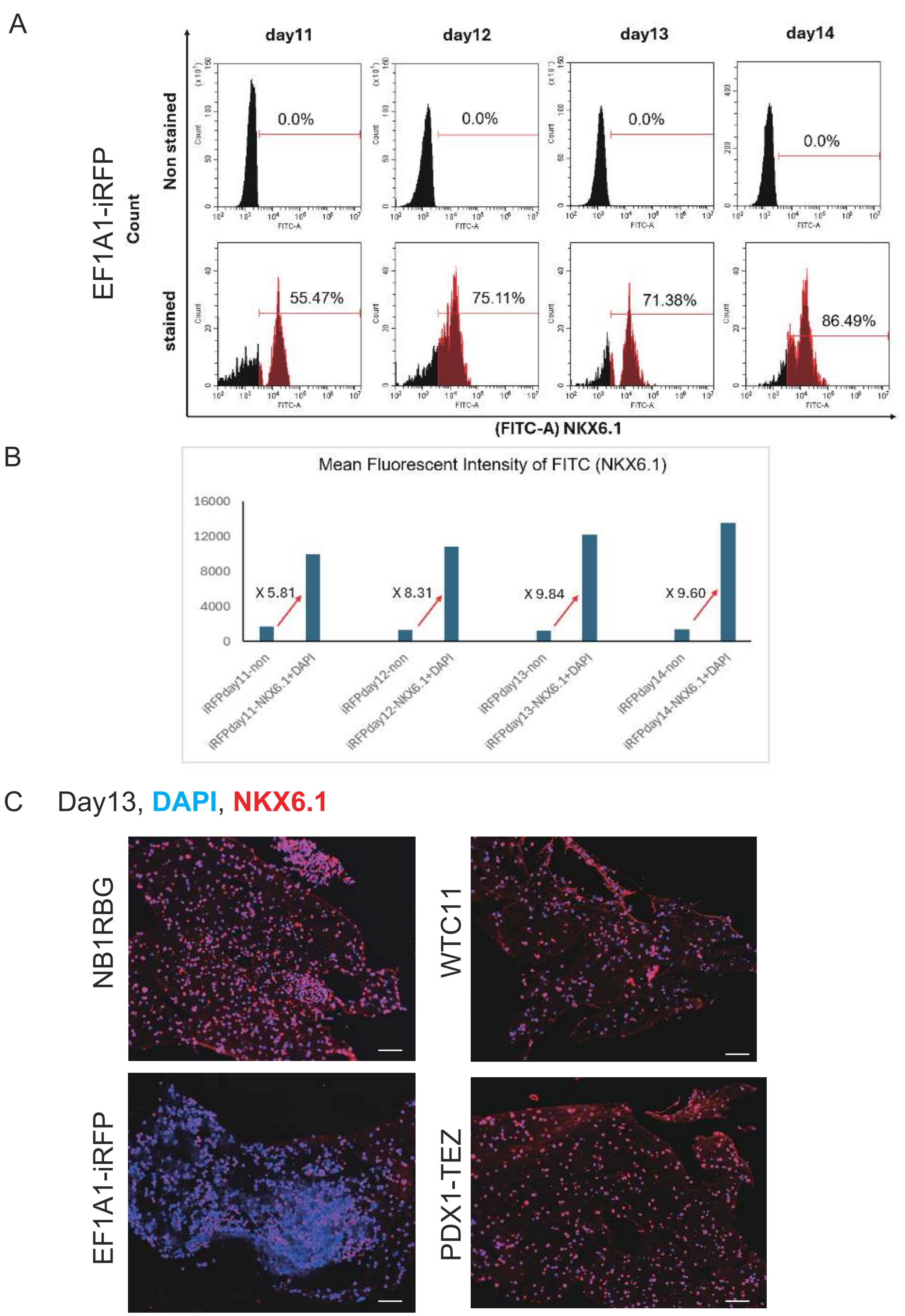


Figure S4 Reproducible NKX6.1 expression across multiple hiPSC lines

(A) Time-course flow cytometric analysis of NKX6.1 expression in EF1A-iRFP hiPSC line during PP differentiation. Representative histograms at day 11, 12, 13, and 14. (B) Mean fluorescence intensity of NKX6.1 expression in EF1A-iRFP line. The "non" bar represents the non-stained control sample, which serves as the negative baseline for MFI measurement. (C) Immunohistochemistry of NKX6.1 in differentiated EEF1A-iRFP, WTC11, NB1RGB, and PDX1-tdTomato lines. Nuclei were counterstained with DAPI (blue).

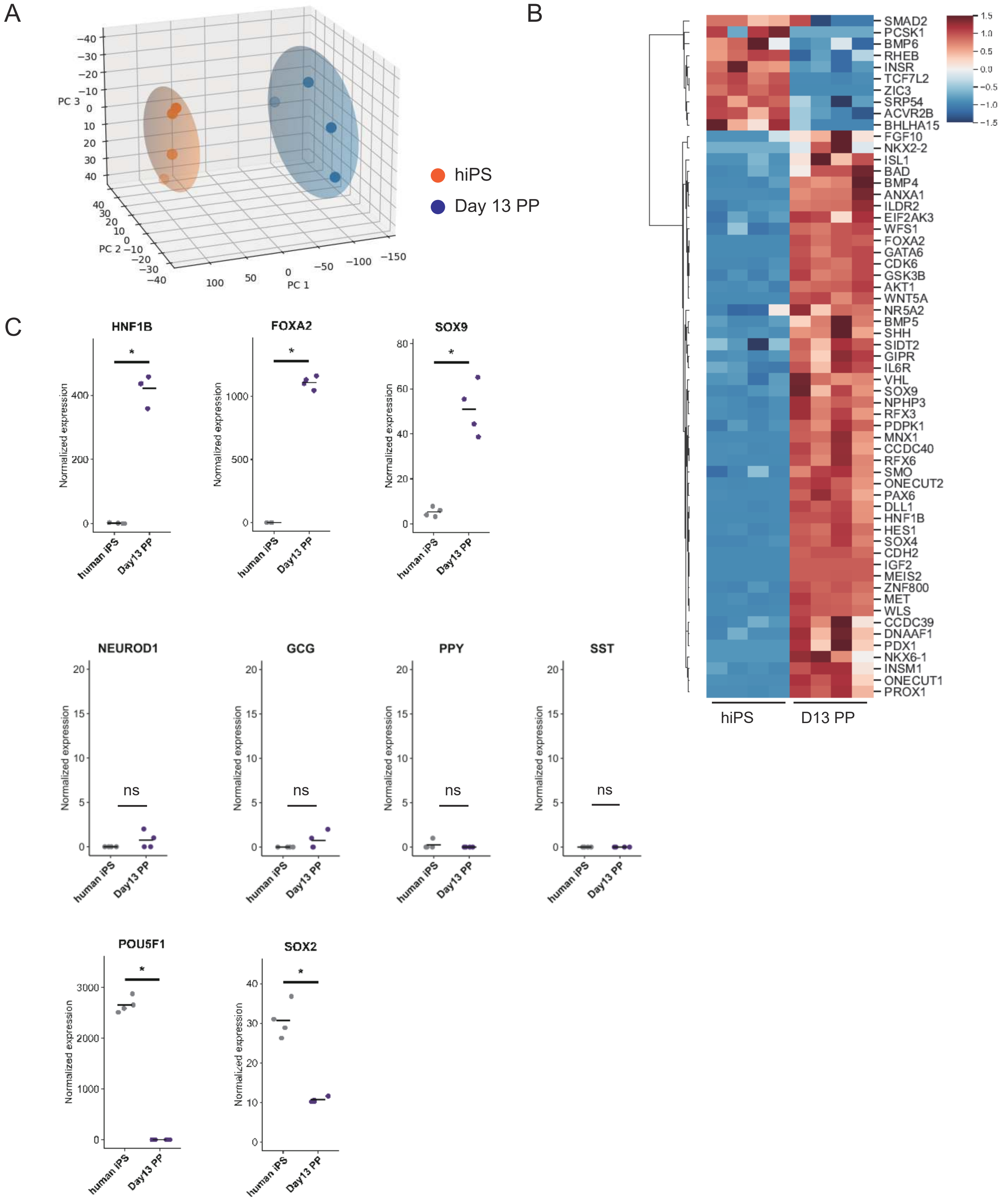


Figure S5 Transcriptome analysis confirms pancreatic progenitor identity after differentiation

(A) Principal component analysis plot of EEF1A-iRFP670 line before (n = 4) and after (n = 4) differentiation. (B) Heatmap showing differential expression of pancreatic development-associated genes between before (n = 4) and after (n = 4) differentiation. (C) Normalized expressions of definitive endoderm markers (*HNF1B*, *FOXA2*, *SOX9*), pancreatic endocrine markers (*NEUROD1*, *GCG, PPY, SST*), pluripotency markers (*POU5F1* and *SOX2*) before (n = 4) and after (n = 4) the differentiation. *$p < 0.05$, ns: not significant.

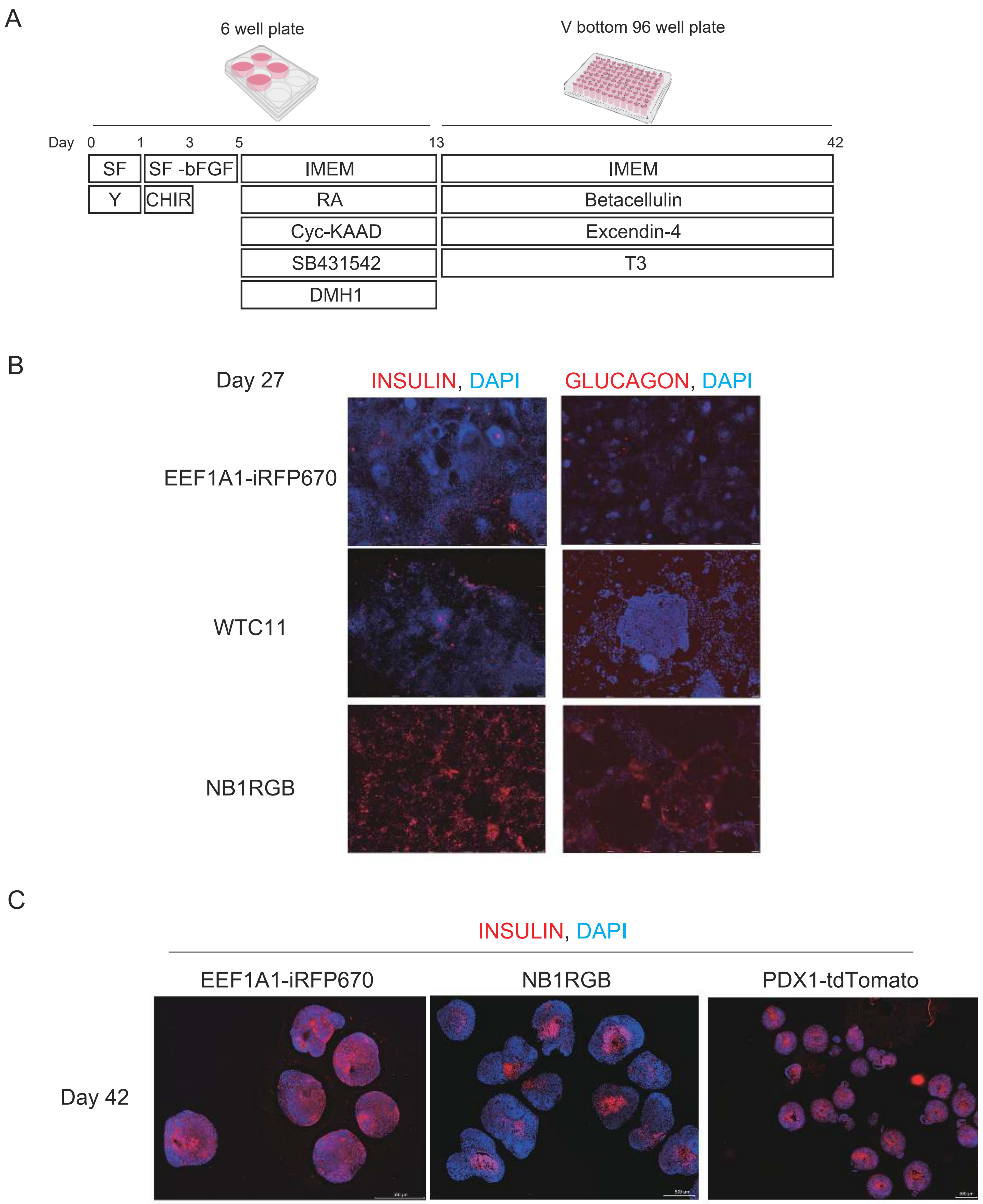


Figure S6 Pancreatic hormone expression in multiple hiPSC lines after extended culture

(A) Schema of the differentiation protocol for differentiating pancreatic endocrine cells from human iPSC lines. (B) Immunostaining of human INSULIN and GLUCAGON on sections of aggregates at day 27. (C) Immunostaining of human INSULIN on sections of aggregates at day 42.

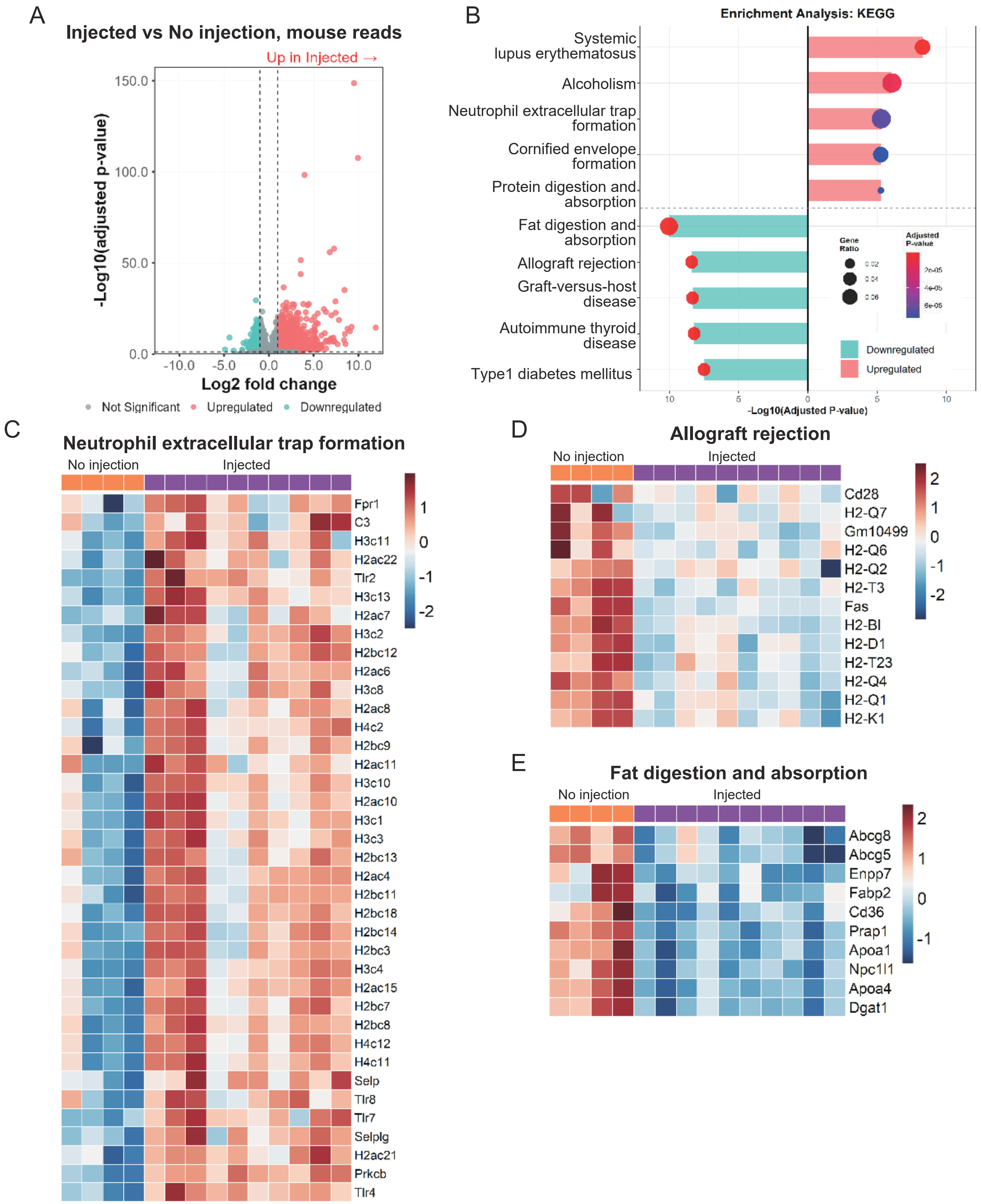


Figure S7 RNA-seq analysis of host response to intraplacental injection

(A) Volcano plot between injected vs no-injection *Pdx1*$^{-/-}$ embryos at E18.5. Red and blue dots indicate significantly up- and down-regulated genes (|log2FC| > 1, adjusted $p$ < 0.05). (B) KEGG pathway enrichment analysis. (C-E) Heatmaps of selected enriched pathways: neutrophil extracellular trap formation (C), allograft rejection (D), and fat digestion and absorption (E). Expression values are z-score normalized. n = 4 for no-injection, n = 10 for injected samples. Statistical analysis by DESeq2 with Benjamini-Hochberg correction.

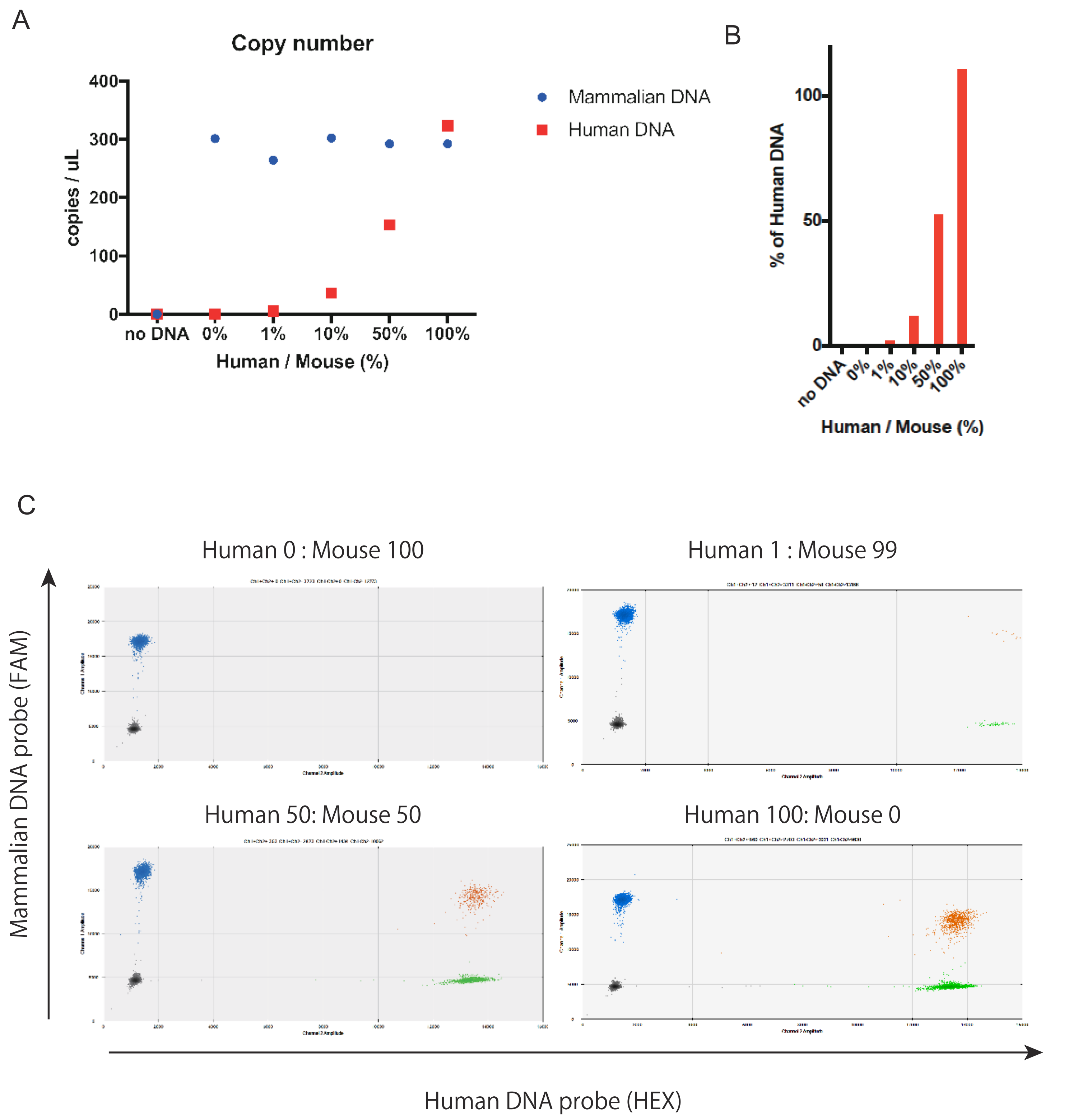


### Figure S8 Validation of human DNA detection specificity in mixed samples by ddPCR

(A) Quantification of total mammalian (blue circles) and human (red squares) DNA in mixed samples with varying human/mouse DNA ratios. (B) Human DNA content calculated as percentage of total mammalian DNA. (C) Representative 2D scatter plots from ddPCR analysis showing distinct populations of positive droplets in samples containing 0% (left-top), 1% (right-top), 50% (left-bottom), and 100% (right-bottom) human DNA.

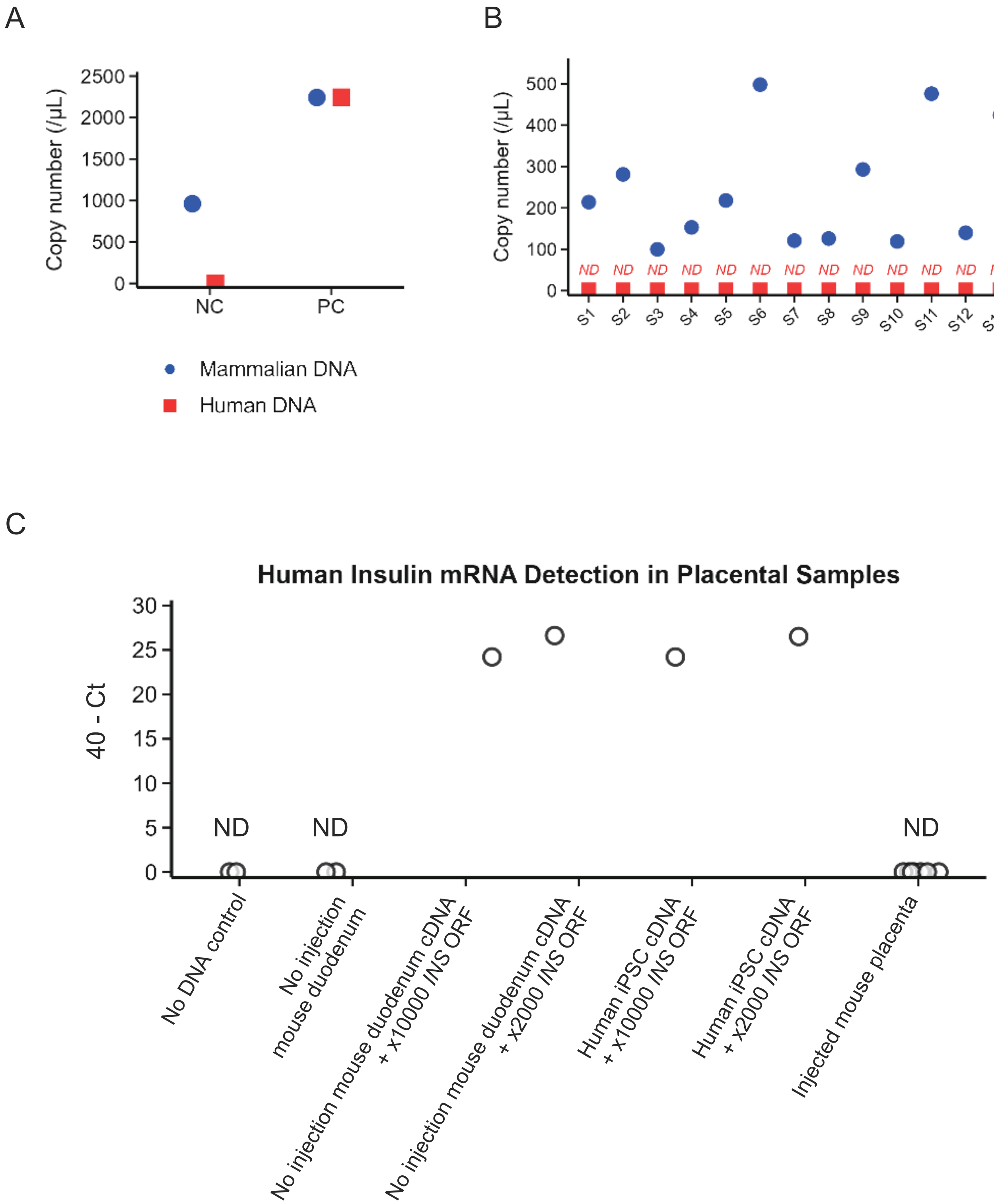


Figure S9 Placental tissue did not retain human cell and ectopic insulin production.
(A) Negative (NC; mouse genomic DNA) and positive (PC; human genomic DNA) controls demonstrating the performance and specificity of the ddPCR assay used for chimerism analysis. (B) ddPCR analysis of individual placental tissues (n = 13, labeled S1-S13) collected at E18.5 from embryos that received intraplacental injections at E9.5. Human DNA was not detected (ND; not detected) in any sample. (C) RT-qPCR analysis for human *INSULIN* (*INS*) in placental tissue from the same individuals in panel (B). No *INS* expression was detected in the transplanted placenta, similar to negative controls (no template, uninjected mouse duodenum). Strong signals (40 - Ct = 25-28) were detected in positive controls consisting of mouse cDNA mixed with a human INS expression plasmid.

Supplemental Table 1

| WT mating | | | |
|---|---|---|---|
| Concentration | Injection | Birth | Birth rate |
| cells/embryo/μl | n of embryos | n of pups | % |
| $2x10^5$ | 27 | 10 | 37.037 |
| $1x10^5$ | 8 | 3 | 37.500 |
| $5x10^4$ | 15 | 2 | 13.333 |
| $2.5x10^4$ | 32 | 12 | 37.500 |
| $1x10^4$ | 15 | 13 | 86.667 |
| $5x10^3$ | 16 | 13 | 81.250 |
| $2.5x10^3$ | 7 | 7 | 100.000 |
| 0 (PBS) | 28 | 25 | 89.286 |

Supplemental Table 2

| *Pdx1*$^{+/-}$ mating | | | | | |
|---|---|---|---|---|---|
| Injection | Birth | Birth rate | Genotype | | |
| n of embryos | n of pups | % of injection | | n of pups | % of pups |
| 82 | 67 | 81.707 | *Pdx1*$^{-/-}$ | 14 | 20.896 |
| | | | *Pdx1*$^{+/-}$ | 38 | 56.716 |
| | | | Wild type | 15 | 22.388 |

Supplemental Table 3

| Antigen | Host | Conjugation | Antibody used<br>Provider | Cat# | Application | Dilution |
|---|---|---|---|---|---|---|
| HLA-Class 1 | Mouse | Biotin | abcam | ab110665 | FACS | 2 µL/million cells /100 µL |
| mCD45.2 | Mouse | FITC | BioLegend | 109806 | FACS | 2 µL/million cells /100 µL |
| mCD31 | Rat | PE | BD bioscience | 553373 | FACS | 2 µL/million cells /100 µL |
| INS | Guinea Pig | none | abcam | ab7842 | IHC | 1:200 |
| Pig IgG | Goat | Alexa Fluor® 488 | Invitorogen | A11073 | IHC | 1:500 |
| OCT3/4 | Mouse | none | Santa Cruz | sc-5279 | IHC | 0.4 µg/mL |
| NANOG | Rabbit | none | Reprocell | RCAB004P-F | IHC | 0.5 µg/mL |
| PAX6 | Rabbit | none | MBL | PD022 | IHC | 1:400 |
| TUJ1 | Mouse | none | R&D Systems | MAB1195 | IHC | 1 µg/mL |
| HAND1 | Goat | none | R&D Systems | AF3168 | IHC | 1 µg/mL |
| SMA | Mouse | none | R&D Systems | MAB1420 | IHC | 1 µg/mL |
| SOX17 | Goat | none | R&D Systems | AF1924 | IHC | 0.4 µg/mL |
| AFP | Mouse | none | R&D Systems | MAB1368 | IHC | 1 µg/mL |
| PDX1 | Goat | none | R&D Systems | AF2419 | IHC | 1:200 |
| CD117 | Mouse | PE | BioLegend | 313203 | FACS | 5 µL/million cells /100 µL |
| CXCR4 | Mouse | APC | BioLegend | 306509 | FACS | 5 µL/million cells /100 µL |
| NKX6.1 | Mouse | Alexa Fluor® 488 | BD bioscience | 567608 | FACS | 5 µL/million cells /100 µL |
| NKX6.1 | Goat | none | R&D Systems | AF5857 | IHC | 1:100 |
| Goat IgG | Donkey | Alexa Fluor® 594 | Invitrogen | A-11058 | IHC | 1:500 |

Supplemental Table 4

| List of Primers for RT-qPCR | | | |
|---|---|---|---|
| S.no. | Gene | Forward (5' to 3') | Reverse(5' to 3') |
| 1 | RPLP0 | GAAACTCTGCATTCTCGCTTCC | ACTCGTTTGTACCCGTTGATGA |
| 2 | NANOG | AAAGAGGTCTCGTATTTGCTGC | GAAACACTCGGTGAAATCAGGG |
| 3 | SOX17 | GCATGACTCCGGTGTGAATCT | TCACACGTCAGGATAGTTGCAGT |
| 4 | HNF6 | TGTGGAAGTGGCTGCAGGA | TGTGAAGACCAACCTGGGCT |
| 5 | PDX1 | CGGAACTTTCTATTTAGGATGTGG | AAGATGTGAAGGTCATACTGGCTC |
| 6 | PTF1A | GAAGGTCATCATCTGCCATCG | GGCCATAATCAGGGTCGCT |
| 7 | NGN3 | GGCTGTGGGTGCTAAGGGTAAG | CAGGGAGAAGCAGAAGGAACAA |
| 8 | NKX6.1 | GGGCTCGTTTGGCCTATTCGTT | CCACTTGGTCCGGCGGTTCT |
| 9 | INS | ACGAGGCTTCTTCTACACACCC | TCCACAATGCCACGCTTCTGCA |
